# Structure of the human TWIK-2 potassium channel and its inhibition by pimozide

**DOI:** 10.1101/2025.02.24.639991

**Authors:** Nandish K. Khanra, Chongyuan Wang, Bryce D. Delgado, Stephen B. Long

## Abstract

The potassium channel TWIK-2 is crucial for ATP-induced activation of the NLRP3 inflammasome in macrophages. The channel is a member of the two-pore domain potassium (K2P) channel superfamily and an emerging therapeutic target to mitigate severe inflammatory injury involving NLRP3 activation. We report the cryo-EM structure of human TWIK-2. In comparison to other K2P channels, the structure reveals a unique ‘up’ conformation of Tyr111 in the selectivity filter and a SF1-P1 pocket behind the filter that could serve as a binding site for channel modulators. Density for acyl chains is present in fenestrations within the transmembrane region that connect the central cavity of the pore to the lipid membrane. Limited pharmacological tools are available for TWIK-2 despite its importance as a drug target. We show that the small molecule pimozide inhibits TWIK-2 and determine a structure of the channel with pimozide. Pimozide displaces the acyl chains and binds below the selectivity filter to block ion conduction. The drug may access its binding site via the membrane, suggesting that other hydrophobic small molecules could have utility for inhibiting TWIK-2. The work defines the structure of TWIK-2 and provides a structural foundation for development of specific inhibitors with potential utility as anti-inflammatory drugs.

**Significance Statement:** The TWIK-2 potassium channel is a member of the two-pore domain potassium (K2P) channel superfamily and a potential therapeutic target to control severe inflammatory injury involving the NLRP3 inflammasome. We report the cryo-EM structure of the human TWIK-2 channel at 2.85 Å resolution, revealing differences in comparison to other K2P channels. We identify that pimozide, an FDA-approved drug for Tourette syndrome, inhibits TWIK-2. A cryo-EM structure of TWIK-2 in complex with pimozide identifies its binding location and mechanism of inhibition. The work provides a structural foundation for development of specific TWIK-2 inhibitors that have potential therapeutic utility for inflammatory diseases involving NLRP3 activation.

## Introduction

Potassium channels have diverse biological functions ranging from propagating action potentials, to controlling the resting membrane potential of cells, to involvement in immune function. Many potassium channels function at the plasma membrane to control potassium efflux from cells, be that in neurons or other cell types, due to the higher potassium concentration within cells than surrounding them. Certain potassium channels function in intracellular organelles, as is the case for TWIK-2, which has been shown to be active in lysosomes (1-6). TWIK-2 also functions at the plasma membrane under certain circumstances. For example, TWIK-2 is one of the main channels responsible for the potassium efflux from macrophages that is necessary to activate the NLRP3 inflammasome (7–11). TWIK-2 is distinct from other potassium channels both functionally and in primary sequence, and its structure has not been determined.

TWIK-2 is a member of the two-pore domain (K2P) potassium channel superfamily (6, 12, 13). Many potassium channels comprise four identical subunits that each contribute one pore-forming domain to constitute the complete channel. The K2P nomenclature of the superfamily is derived from the presence of two pore-forming domains (PD1 and PD2) within each protomer, such that the channel is formed by a dimer of subunits (14, 15). There are fifteen members of the K2P superfamily in humans (16, 17). The channels are expressed widely and function in diverse physiological processes ranging from modulating cardiac function to intra-ocular pressure regulation, to sleep duration, to lung injury responses (18–20). Members of the superfamily are regulated by various physical and chemical stimuli such as pressure, temperature, extracellular and intracellular pH, lipids, and phosphorylation, and some are sensitive to volatile anesthetics and antidepressants (16, 17, 21). Structures of several K2P channels revealed that the transmembrane-spanning region of the pore has a similar architecture to that of tetrameric potassium channels (22–28). These structures also divulged distinct structural features of K2P superfamily members, which include an ‘extracellular cap’ domain positioned above the selectivity filter, lateral ‘fenestrations’ in the transmembrane regions that form openings between the pore and the hydrophobic core of the lipid bilayer, and a lower ‘X-gate’ region within the pore of certain members that functions as a gate to control ion flow.

The K2P superfamily can be divided into six subgroups, delineated based on amino acid sequence similarity and functional properties: TWIK, TREK, TALK, TASK, THIK, and TRESK (16, 17). The TWIK (tandem of pore domain in a weak inward rectifying K^+^ channel) family has two members in humans, TWIK-1 and TWIK-2, but these proteins share only 44.5 % amino acid sequence identity (6, 12–14). The structure of TWIK-1, which was one of the first to be determined for K2P channels (22), defined many of the structural features found in the K2P superfamily. TWIK-1 and TWIK-2 exhibit electrophysiological and physiological differences, suggesting that TWIK-2 may be structurally distinct. For example, the activity of TWIK-1 is controlled by the pH on the extracellular side of the channel, for which a structural basis is known (29, 30), whereas TWIK-2 is not effected by external pH (12). The single channel conductance of TWIK-2 is low in comparison to other K2Ps (13). TWIK-1 and TWIK-2 have distinct expression profiles and function in different physiological processes. TWIK-2 is expressed in the eye, lungs, stomach, spleen, vascular system, and embryo (6, 13, 21, 31). TWIK-1 is abundantly expressed in the heart and brain and to a lesser extent in the placenta, lung, liver, and kidney (14).

TWIK-2 is the only K2P channel observed to exhibit inactivation, wherein currents through the channel decrease over time (13). Inactivation of TWIK-2 occurs under physiological (∼ 5 mM) concentrations of extracellular K^+^, whereas higher (∼ 150 mM) extracellular K^+^ concentrations reveal only modest inactivation (12, 13). Inactivation is observed for outward K^+^ currents, but the channel does not demonstrate inactivation for inward currents (13).

The importance of TWIK-2 in the activation of the NLRP3 inflammasome was recently discovered (7, 32). The NLRP3 inflammasome is a multi-protein complex that plays critical roles in innate immunity and inflammatory signaling (33, 34). It has been reported to play key roles in diverse diseases including sepsis-induced lung injury, cancer, excessive inflammatory response in COVID-19, diabetes, and obesity (33–37). The NLRP3 inflammasome also been shown to be involved in neuroinflammation in response to the accumulation of amyloid-β or ⍺-synuclein in neurodegenerative diseases (38, 39). Prior to activation of the NLRP3 inflammasome in macrophages, TWIK-2 is localized to endosomal compartments (1, 8). The presence of extracellular ATP induces endosomal trafficking of TWIK-2 to the plasma membrane (8). This re-localization allows K^+^ efflux and a reduction of the cytosolic concentration of K^+^, which is necessary for the assembly of the NLRP3 inflammasome (8, 40). Small molecules have been shown to inhibit TWIK-2 and reduce NLRP3 inflammasome activation, but these compounds have modest potency against TWIK-2 (41, 42). Three-dimensional structures of TWIK-2 alone and in complex with inhibitors would be helpful for inhibitor optimization.

We report the cryo-EM structure of human TWIK-2 at 2.85 Å resolution. Comparisons with the structures of other K2P channels reveal similarities in overall architecture and distinctions within the pore, selectivity filter, and extracellular cap domain. Through functional analysis using a reconstituted system, we demonstrate that purified TWIK-2 catalyzes K^+^ flux. Using this system, we show that the small molecule pimozide, which is an FDA-approved drug used for treatment of severe Tourette’s syndrome (43, 44), inhibits TWIK-2. A cryo-EM structure of TWIK-2 in complex with pimozide identifies the location of drug binding and its mechanism of action.

## Results

### Overall Structure of TWIK-2

Full-length human TWIK-2 was expressed in mammalian cells and purified using mild detergents at neutral pH for cryo-EM studies (Materials and Methods, Fig. S1). We determined its cryo-EM structure by single-particle methods to 2.85 Å resolution (Fig. 1 and Figs. S2-3 and Table S1). TWIK-2 forms a symmetric dimer, with the ion pore through the transmembrane domain located along the channel’s two-fold (C2) rotational axis of symmetry (Figs. 1*C* and 1*D*). The overall architecture of the channel resembles that of other K2P channels. An extracellular cap domain is positioned above the transmembrane-spanning ion conduction pore. The pore comprises a selectivity filter region and a central cavity that would be exposed to the intracellular solution. Each subunit contains two extracellular helices (E1 and E2), four transmembrane helices (M1-M4), two pore helices (P1 and P2), and two selectivity-filter regions (SF1 and SF2). A single ion conduction pore containing the ion selectivity filter is formed from the dimeric assembly of the subunits. Potassium ions flowing through the channel from the intracellular to the extracellular side would enter through the wide intracellular opening, pass through the central cavity, selectivity filter, and emerge on the extracellular side below the cap domain, where they could reach the extracellular solution through bifurcated gaps between the cap and the selectivity filter (Fig. 1*D*). Like other K2P channels visualized to date (17), the channel has a domain-swapped arrangement wherein the transmembrane M1 helix and the extracellular E1 helix of one subunit are positioned next to the other subunit rather than its own (Fig. 1). TWIK-1 was previously thought not to have such an arrangement from the initial crystal structure, but it was also shown to have this domain-swapped architecture (22, 30). The cryo-EM structure of TWIK-2 has a well-defined density for the majority of the channel (Figs. S3*F*-*L*). Only the terminal ends (amino acids 1-6 and 272-313) and the loops connecting M2 to M3 (amino acids 152-165) and part of E2 to P1 (amino acids 79-90) are disordered. The disordered part of the E2 to P1 loop contains two glycosylation sites (Asn79 and Asn85). The glycosylation of these residues has been shown to be important for localization of TWIK-2 in lysosomes (1).

**Fig. 1.**
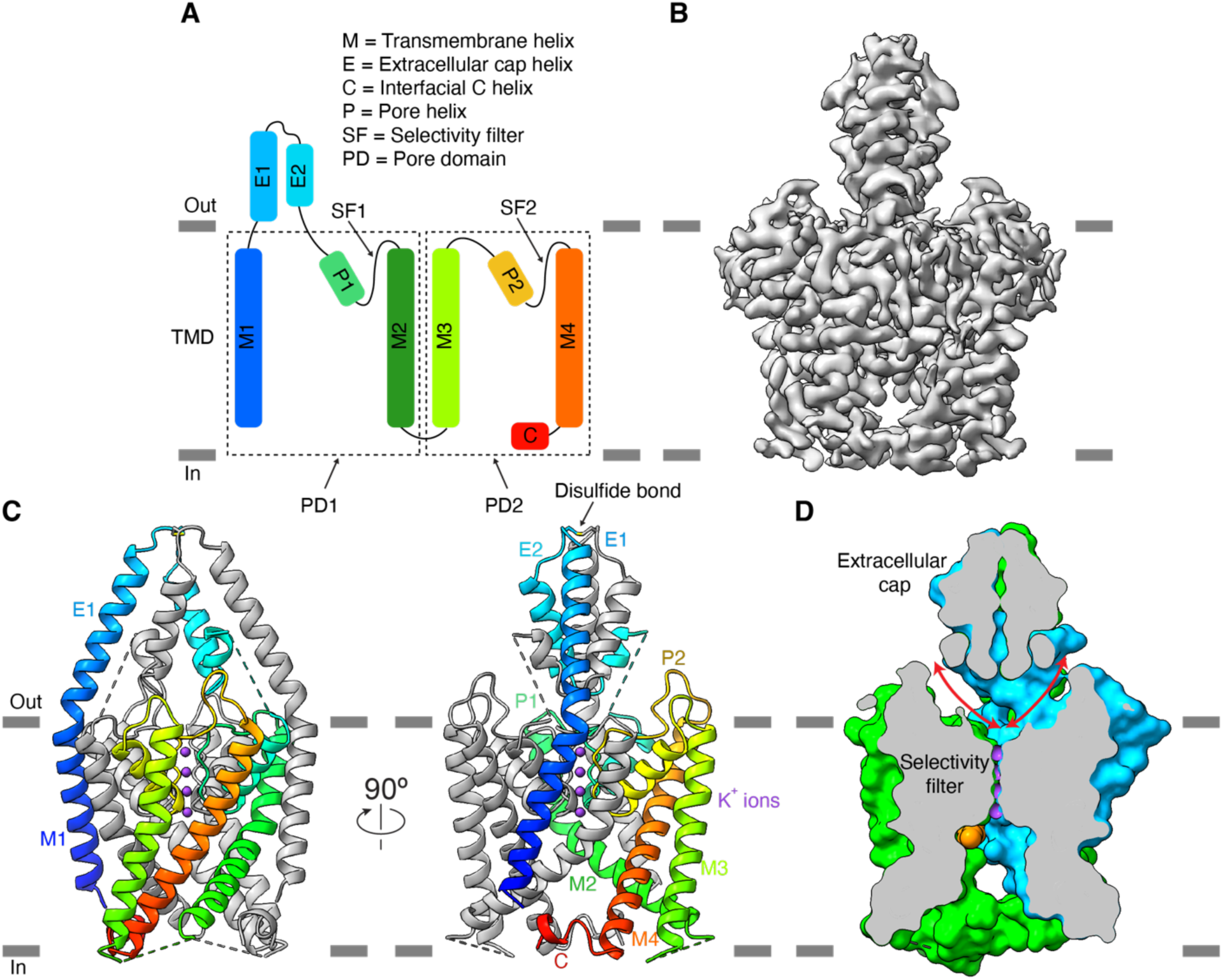
Overall structure of TWIK-2. *(A)* Secondary structure schematic of one subunit, colored blue-to-red from the N- to C-terminus. Horizontal bars indicate approximate membrane boundaries. The dashed boxes represent the two pore-forming domains of the TWIK-2 gene (PD1 and PD2). *(B)* Cryo-EM density. *(C)* Architecture of the channel, shown from two perspectives. One subunit is grey, and the other is colored as in *(A)*. Dashed lines indicate disordered loop regions. *(D)* Cutaway view of the molecular surface of the channel from the side. Arrows indicate the bifurcated extracellular ion passageways. Potassium ions are shown as purple spheres. An acyl chain is orange (sphere representation).

### Selectivity filter

The selectivity filter is located near the extracellular side of the pore (Fig. 1). Owing to the similarities between the regions of each subunit that contribute to the filter (SF1 and SF2, and P1 and P2), the selectivity filter has pseudo four-fold symmetry (Fig. 2). The protein sample used for the structure contained 150 mM K^+^, a concentration that was chosen to mimic a typical cellular concentration of K^+^. Cryo-EM density consistent with K^+^ is present at four sites (S1 through S4) in the filter, as observed in other K^+^ channels (Fig. 2*A*) (45, 46). S3 has the strongest density, but this could be due to the higher local resolution of this region (Fig. S3). As first observed for the prototypic potassium channel KcsA (45), K^+^ in sites S1-S3 are coordinated exclusively by backbone carbonyl oxygen atoms (Figs. 2*A*-*B*). As in KcsA, the potassium ion in site S4 is coordinated by both backbone carbonyl oxygens and the hydroxyl moieties of threonine residues – in TWIK-2, these are contributed by Thr106 and Thr214 from PD1 and PD2, respectively (Figs. 2*A*-*B*). In the TWIK-2 structure, ion coordination is four-fold symmetric – the backbone conformations of SF1 and SF2 and the side chain positions of Thr106 and Thr214 adopt the same conformations within the error of the coordinates (Figs. 2*A-B*). These conformations correspond to conductive conformations observed in other K2P and K^+^ channel structures (Fig. 2*C*) (45, 46). The asymmetry of the amino acid sequences comprising SF1 and SF2 of TWIK-2 are accommodated by different packing of their side chain residues with the surrounding regions (Fig. 2*E*) to engender the four-fold symmetric nature of ion coordination within the filter. The presence of density at S1-S4 and a backbone structure analogous to the conductive conformation of the selectivity filter of other K^+^ channels suggest that the filter represents an ion-conductive state.

**Fig. 2.**
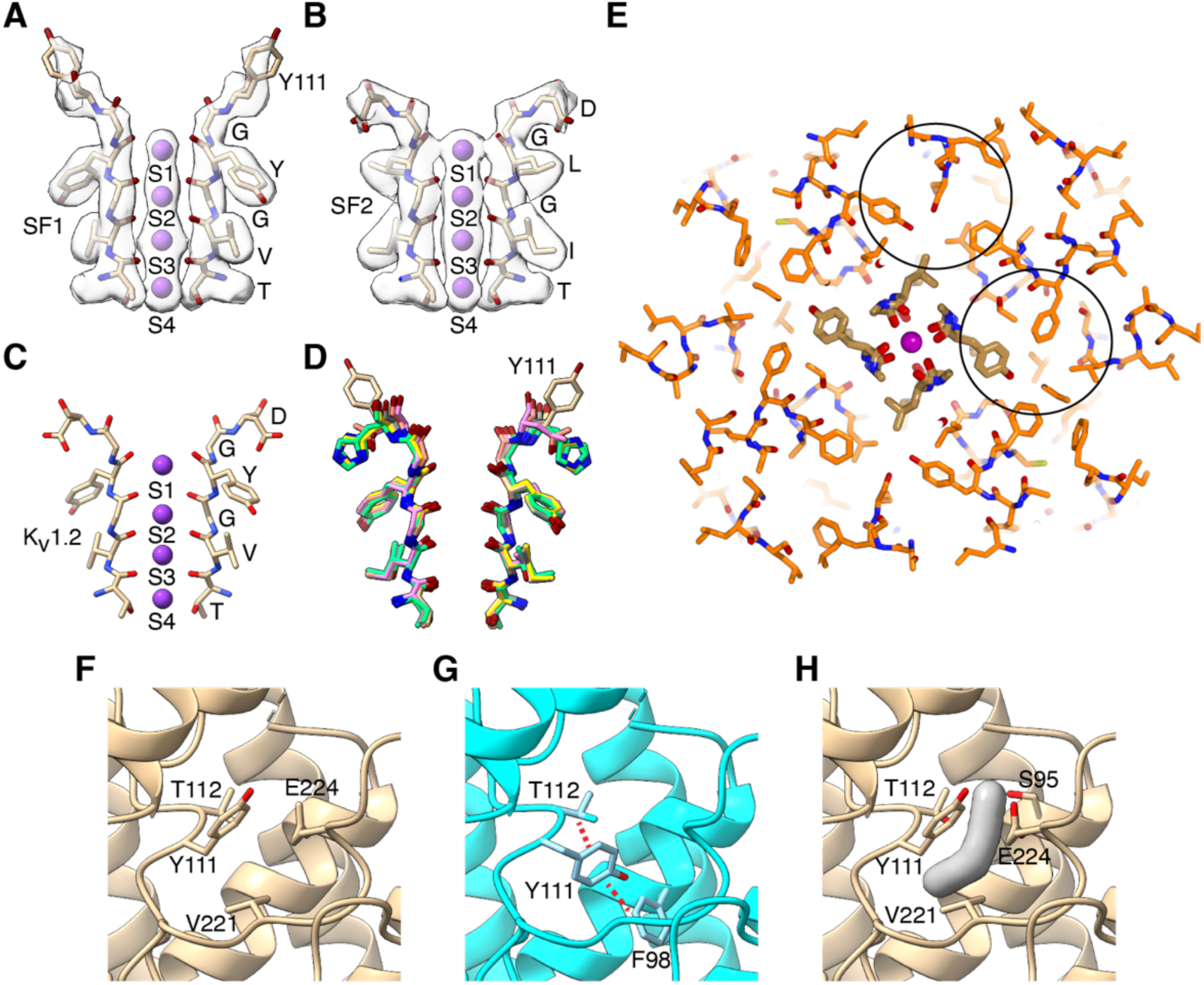
The selectivity filter of TWIK-2. *(A)* Selectivity filter 1 (SF1) region. The atomic model is drawn as sticks, with potassium ions represented as spheres. Cryo-EM density is shown as a semitransparent surface. *(B)* Selectivity filter 2 (SF2) region depicted as in *(A)*. *(C)* The selectivity filter of Kv1.2 (PDB ID 2R9R). *(D)* Comparison of SF1 from TWIK-2 with other K2P channels. The position of Tyr111 is noted. Shown are: TWIK-2 (tan), TWIK-1 (green, 3UKM), TREK1 (grey, 8DE8), TREK2 (salmon, 4BW5), TRAAK (violet, 3UM7), TASK1 (blue, 6RV4), and TASK3 (gold, 8K1J). *(E)* Cutaway view of the selectivity filter from the top, showing pseudo four-fold symmetry and the packing of surrounding residues. The atomic model is depicted as sticks, with carbon atoms of SF1 and SF2 in yellow and the remainder in orange. Oxygen and nitrogen atoms are red and blue, respectively. *(F)* Region surrounding Tyr111. Tyr111 is shown in its observed conformation and depicted as sticks. *(G)* Hypothetical modelling of Tyr111 in a ‘down’ conformation, showing steric clashes as red dashed lines. (H) The SF1-P1 pocket, depicted as a grey surface.

### Tyr111

Tyr111 in position six of the SF1 sequence (TVGYG<u>Y</u>) deviates from the canonical K^+^ channel filter sequence and most other K2P channels. In KcsA, Kv1.2 and other tetrameric K^+^ channels, the corresponding amino acid is usually an aspartic acid. In K2P channels, this amino acid varies but is typically His, Asn, or Met (Fig. S4). In only TWIK-2 and human TRESK is a tyrosine present at that position. In the structure of TWIK-2, Tyr111 adopts an ‘up’ conformation (Figs. 2*A* and *F*). In the structures of tetrameric potassium channels and other K2P channels that have been determined, the amino acid at position six is typically in a ‘down’ conformation, where it packs behind the selectivity filter (Fig. 2*D* and Fig. S6). The corresponding amino acid in TWIK-1, His122, has been shown to be responsible for that channel’s sensitivity to extracellular pH (29, 30). Dramatic conformational changes occur in the selectivity filter of TWIK-1 due to pH changes – at neutral pH, the filter adopts the canonical conductive conformation, and His122 is down, while at acidic pH, His122 adopts an ‘up’ conformation and the filter collapses into a non-conductive state (30). The presence of tyrosine at this position in TWIK-2 probably explains the pH insensitivity of the channel relative to TWIK-1. In the structure of TWIK-2, there is a pocket located behind the filter (the ‘SF1-P1 pocket’) at the interface between SF1 and the P1 helix (Fig. 2*H*). The SF1-P1 pocket corresponds approximately to the position occupied in other channels by the ‘down’ conformation of amino acid six (Fig. S6). Modeling of Tyr111 in a ‘down’ rotamer produces molecular clashes with surrounding amino acids (Fig. 2*G*), but these could conceivably be eliminated by minor repositioning of these and nearby residues (Phe98, Thr112, Val221, and Lys235). Therefore, it seems plausible that Tyr111 could adopt a down position under certain circumstances. We attempted to isolate different conformations of the selectivity filter from focused 3D classifications of the single particle dataset; these efforts yielded only the consensus structure reported here with Tyr111 in the up position. A pocket behind the selectivity filter of the K2P channel TREK-1, in a different position (the P1-M4 interface), is a site for control of channel function by small molecules, whereby the presence of the small molecule stabilizes the conductive conformation of the selectivity filter (47). The SF1-P1 pocket observed in the structure of TWIK-2 could potentially be targeted for the development of small molecules that may modulate channel activity.

### The pore

Below the selectivity filter, the pore widens to form a central cavity, the diameter of which is large enough to accommodate hydrated potassium ions (Figs. 3*A*-*B*). The cavity is lined by the ‘inner’ helices of the channel (M2 and M4). The transmembrane helices (M1-M4) adopt conformations similar to those observed in the structure of TWIK-1 (Figs. S5*A*-*B*). While the structures of some K2P channels have revealed a constriction near the cytosolic side of their pores, referred to as an ‘X gate’ (28), the TWIK-2 structure does not have this constriction, suggesting that ion flow through the channel is not controlled (‘gated’) by movements of the transmembrane helices. The side chain of Met135 on PD1 marks the narrowest region within the central cavity (Figs. 3*B*-*D*). The corresponding amino acid on PD2 is Val250. The size differential between a methionine and a valine gives this region of the cavity an asymmetric oval-like shape (Fig. 3*D*). Met135 and Val250 correspond to Leu146 and Leu261 in TWIK-1 and likewise constitute the narrowest constriction of the central cavity in that channel (Fig. S5*C*). The cytosolic side of the pore in TWIK-2 is flanked by a C-terminal region of the polypeptide, the ‘C helix’ (Fig. 1*C*). The C helix of TWIK-2 is shorter than in TWIK-1, but generally occupies the same location (Fig. S5*D*-*E*). This helix is a characteristic feature of the TWIK family, observed only in TWIK-1 and TWIK-2 among the K2P channel structures determined to date.

**Fig. 3.**
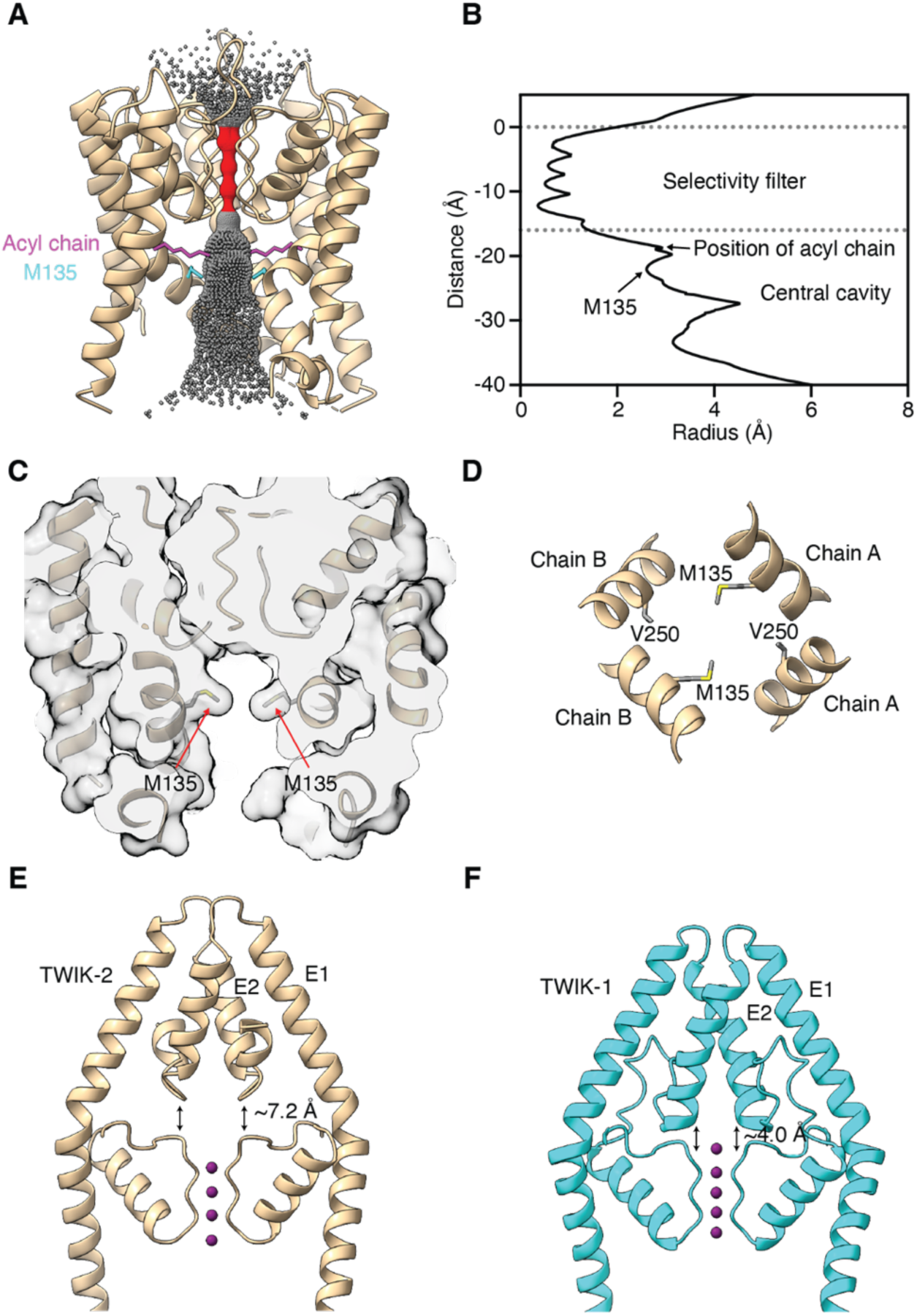
The pore, central cavity, and extracellular cap of TWIK-2. *(A)* Cutaway view of the channel from the side, showing the pore. The pore is depicted by dots representing the distance from its center to the nearest van der Waals contact. The red region represents the selectivity filter in which K^+^ ions become laterally dehydrated. Grey denotes wider regions. Parts of M2 and M4 closest to the viewer are hidden for clarity. Acyl chains (magenta) and Met135 sidechains (blue) are drawn as sticks. *(B)* Dimensions of the pore from *(A)*, depicted as the minimal radial distance to the nearest van der Waals contact. The zero coordinate on the Y-axis corresponds to the carbonyl oxygen of Gly110. *(C)* Met135 residues of chains A and B form a constriction inside the central cavity. A cutaway view of the transmembrane region of the channel is shown. The atomic model is depicted as cartoons and sticks (for Met135) within a semi-transparent molecular surface. *(D)* The Met135 constriction in the central cavity. Portions of the inner helices (M2 and M4) from both subunits are shown with Met135 and Val250 drawn as sticks. Perspective is looking down the pore. *(E)* Extracellular cap of TWIK-2, showing the gap between the selectivity filter and the cap. The distance indicated is measured between the Cα positions of Arg71 and Thr112. *(F)* Comparison with the extracellular cap of TWIK-1 (PDB 7SK0), depicted as in *(E)*. K^+^ ions are shown as spheres; a fifth ion is observed in TWIK-1 between the cap and the filter.

### Acyl densities in fenestrations

Another structural feature of the central cavity is the presence of lateral membrane openings (‘fenestrations’) in the molecular surface of the protein that create windows between the ion pore and the lipid bilayer surrounding the transmembrane region of the channel (Fig. 4*A*). Similar fenestrations have been observed in all K2P channels visualized to date (22–28). The fenestrations are located at interfaces between the two subunits of the channel (Fig. 4*A*). We observe four fenestrations in total, two at each subunit interface: an upper fenestration slightly below the level of the selectivity filter and a lower fenestration located nearer the cytosolic side (Fig. 4*A*). Cryo-EM density is present within each upper fenestration consistent with an acyl chain, which could originate from detergent or lipid molecules (Figs. 4*B-D*). Similar densities have been observed in other K2P channels within their fenestrations and also attributed to acyl/lipid molecules (22, 24, 26, 28, 30). The acyl density is located below SF2, but it does not appear to block the ion permeation pathway (Figs. 4*C*-*D*). Its positioning differs from that observed in the structure of TWIK-1, where the acyl density is positioned directly below the ions in the selectivity filter such that it might impede ion permeation (22). In TWIK-2, the acyl moiety makes van der Waals contacts with Val131, Pro132, and Ser213 of one protomer and Leu246, Met249, and Leu253 of the other protomer (Fig. 4*E*).

**Fig. 4.**
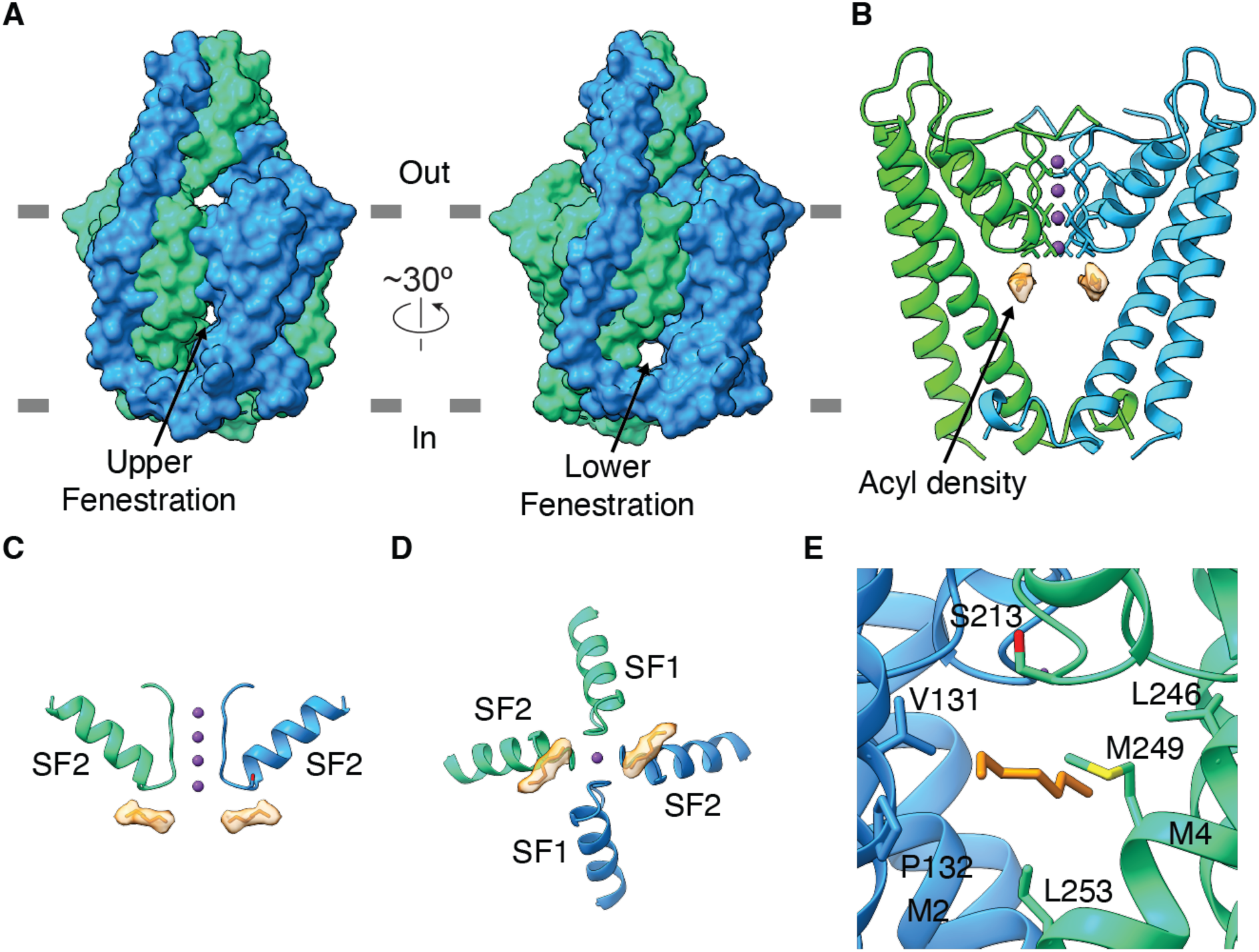
Fenestrations and acyl chain densities in TWIK-2. *(A)* Surface representations of TWIK-2, showing the upper and lower fenestrations. The two subunits of TWIK-2 are shown in blue and green. *(B)* Density for the acyl chains (orange) in the context of a cutaway view of the channel. Density is depicted as semi-transparent surface. The cap and portions of the transmembrane region are hidden for clarity. The densities are in the upper fenestrations. *(C-D)* Depiction of the acyl densities relative to the selectivity filter, shown from the side *(C)* and from below *(D). (E)* Residues surrounding one of the acyl chains. Amino acids within van der Waals contact distance of one of the acyl chains (orange) are depicted as sticks in the context of a cartoon representation of the upper fenestration region. The perspective is from within membrane.

### Extracellular cap domain

The extracellular cap domain is a structural feature, only known to exist in K2P channels, that forms an archway over the extracellular mouth of the pore (22, 23). Ions exiting from the selectivity filter of TWIK-2 would reach the extracellular solution through bifurcated gaps between the cap and the filter (Fig. 1*D*). The cap consists of helices E1 and E2 and intervening loop regions from each subunit. Like most K2P channels, TWIK-2 has a conserved Cys53 in each protomer that forms a disulfide bond with the same residue from the other subunit at the apex of the cap (Fig. 1*C*). Mutation of Cys53 to Ser has been shown to reduce the functional expression of TWIK-2 (13). In comparison to TWIK-1, there is substantially more space between the cap domain and the selectivity filter of TWIK-2 through which hydrated K^+^ ions would pass unobstructed (Figs. 3*E*-*F*). The extra space results from a shorter E2 helix in TWIK-2 in comparison to TWIK-1 (Figs. 3*E*-*F*). In TWIK-1, closure of the gap between the filter and the cap at low pH contributes to the pH dependence of that channel (30). Although the function of TWIK-2 is not regulated by extracellular pH, it seems possible that the cap could have a yet-unknown regulatory function in TWIK-2.

### Purified TWIK-2 is functional and inhibited by pimozide

We sought to assess the function of the purified protein. For this purpose, TWIK-2 was reconstituted into liposomes and studied using a fluorescence-based flux assay analogous to one previously described for TWIK-1 (Fig. 5*A*) (22). The liposomes were filled with 150 mM KCl, whereas the outside buffer contained 150 mM NaCl, creating a K^+^ gradient. Efflux of K^+^ through the channel (out of the liposomes) will produce a negative electric potential within the liposomes that can be used to drive the uptake of protons through an ionophore (CCCP) and quench the fluorescence of a pH indicator (ACMA). A time-dependent decrease in fluorescence indicated K^+^ efflux through the TWIK-2 channel (Figs. 5*B-D*). The assay indicated the selectivity of the channel for K^+^. If Na^+^ were equally permeable, no decrease in fluorescence would be observed. We tested the effect of Ba^2+^, which has been shown to inhibit TWIK-2 using whole-cell patch clamp (13), and found that it inhibits purified TWIK-2 (Fig. 5*B*). We also evaluated the effect of the small molecule ML365, which was previously reported to inhibit TWIK-2 using a cell-based assay (42), and confirmed that this compound inhibits TWIK-2 directly (Fig. 5*C*). From these findings, we conclude that the reconstituted protein recapitulates known properties of TWIK-2. Finally, we assessed the effect of pimozide on the channel. Pimozide is an FDA-approved antipsychotic drug to treat resistant tics and severe symptoms of Tourette syndrome (43, 44). It is an antagonist for some dopamine D_2_-receptors and 5-HT_7_ receptors (43, 48). A previous study suggested that pimozide might inhibit TWIK-2 in the murine lung (49). We evaluated 0.6 and 2 µM concentrations of pimozide and found that pimozide inhibits TWIK-2 in a dose-dependent manner (Fig. 5*D*). The apparent potency of pimozide was greater than that of ML365, which displayed similar levels of inhibition at 10 µM.

**Fig. 5.**
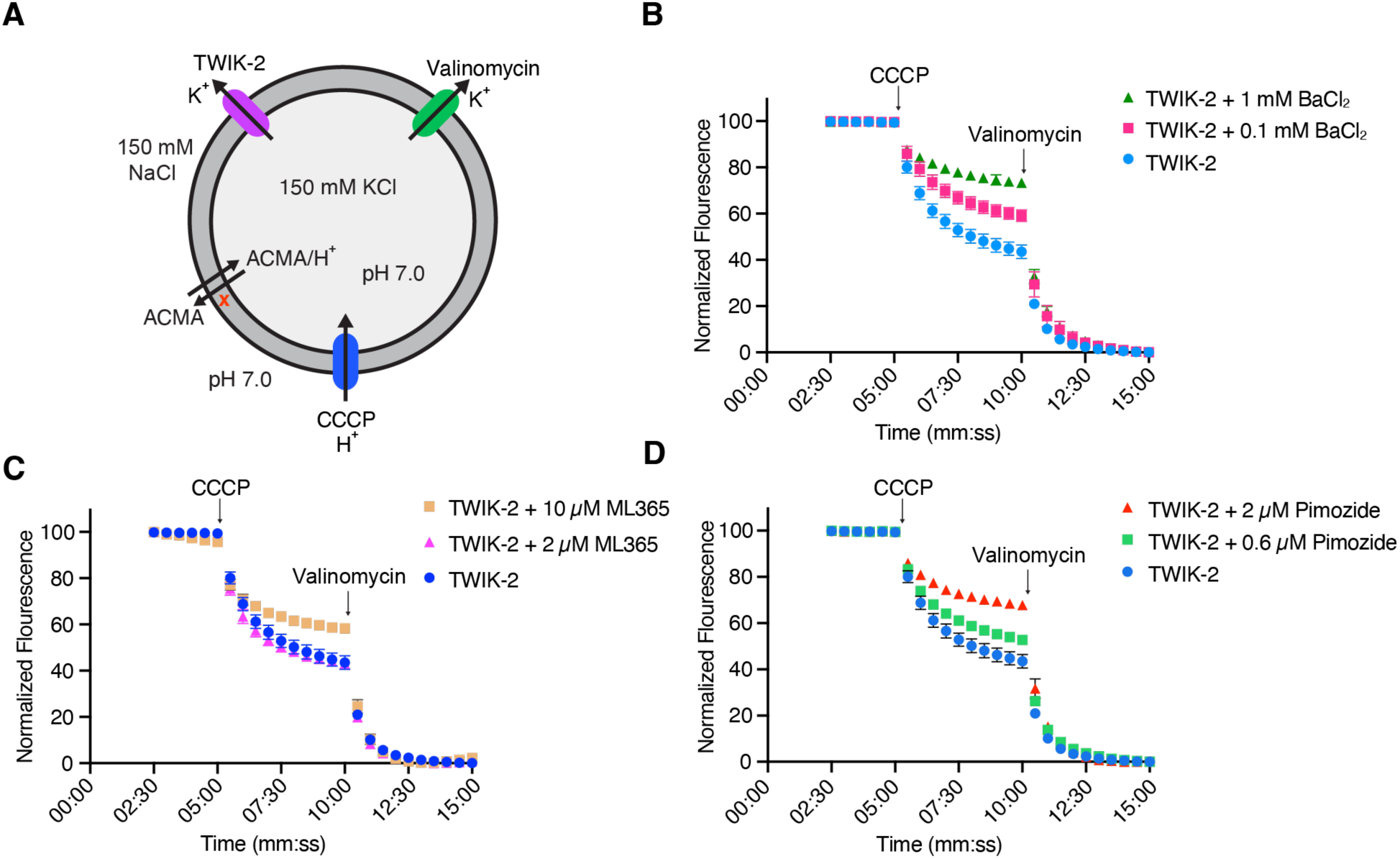
Reconstitution of TWIK-2 channel function and inhibition by pimozide. *(A)* Schematic of the fluorescence-based flux assay. Purified TWIK-2 was reconstituted into liposomes containing 150 mM KCl. These proteoliposomes were diluted into 150 mM NaCl to establish cation gradients. K^+^ efflux through TWIK-2 (out of the proteoliposomes) produces a negative electric potential within the liposomes that drives the uptake of protons through an ionophore (CCCP) and quenches the fluorescence of a pH indicator (ACMA). The K^+^ ionophore valinomycin is added at the end of the experiment to establish a fluorescence baseline and confirm liposome integrity. *(B)* Flux measurements showing that TWIK-2 catalyzes K^+^ efflux. A titration shows that Ba^2+^ (as BaCl_2_) inhibits K^+^ flux through TWIK-2. *(C)* Inhibition of TWIK-2 by ML365. *(D)* Inhibition of TWIK-2 by pimozide. Fluorescence values were normalized by subtracting the value at 15 min (after the valinomycin baseline) and then dividing by the value before CCCP addition. Error bars represent SEM of three independent experiments; some error bars are smaller than data points.

### Pimozide binds below the selectivity filter

To understand the mode by which pimozide inhibits TWIK-2, we determined the cryo-EM structure of TWIK-2 in the presence of pimozide. The structure is determined to 3.2 Å resolution under analogous conditions as the apo structure (150 mM KCl and pH 7.5) (Figs. S7-S9). Strong non-protein density consistent with pimozide is present in the central cavity below the selectivity filter (Fig. 6). The density is located between the selectivity filter and the narrowest region of the central cavity formed by Met135. The shape of the density corresponds to pimozide binding in two equivalent conformations, according to the 2-fold rotational symmetry of the TWIK-2 channel (Fig. 6*C*). We were unable to computationally separate these two conformations using single particle methods. In the structure with pimozide, Met135 adopts two alternative rotamer conformations (Fig. 6*C*), whereas this residue adopts only one rotamer in the absence of pimozide (Figs. S11*A*-*B*). Aside from Met135, the protein conformations in the apo and pimozide structures are indistinguishable (within the limit of the resolution; RMSD = 0.37 Å for Cα positions, Fig. S11*C*). A single pimozide molecule displaces both acyl chains observed in the apo structure. The fluorophenyl and benzimidazole-like moieties on either end of the central piperidine ring of pimozide occupy roughly the same locations as the acyl densities (Figs. 4*C*-*D* and 6*D*-*E*). The piperidine ring binds directly below the selectivity filter, a location in which density is not observed in the absence of pimozide.

**Fig. 6.**
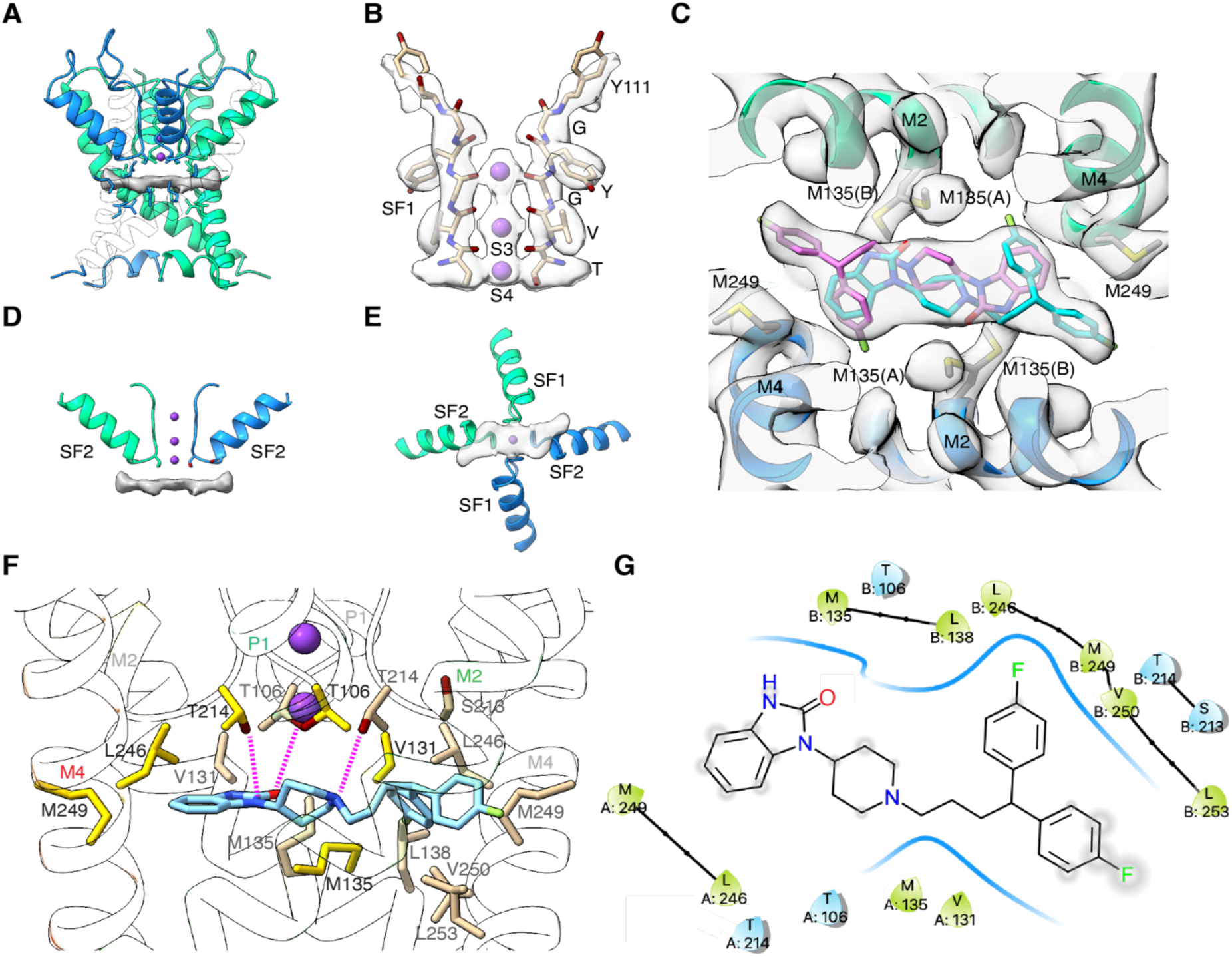
Cryo-EM structure with pimozide. *(A)* The pimozide binding site is below the selectivity filter, spanning the central cavity and upper fenestrations. Density for pimozide is shown in grey surface representation. The transmembrane portion of TWIK-2 is shown in cartoon representation with some regions semi-transparent. *(B)* The selectivity filter (SF1) of the pimozide complex, shown in the same representation as Fig. 2A. Cryo-EM density is drawn in semi-transparent surface representation. *(C)* Cryo-EM density for pimozide. Density for the region in which pimozide binds is shown in semi-transparent surface representation. The density for pimozide corresponds to two equivalent orientations, related by 180*°*, according to the C2 symmetry of the channel. The two pimozide orientations (sticks) are colored separately (with light blue or magenta carbon atoms). Heteroatoms are colored blue for nitrogen, red for oxygen, and green for fluorine. Density for a portion of the surrounding protein highlights the two alternate conformations of Met135 (labeled ‘(A)’ and ‘(B)’) on both chains. *(D-E)* Pimozide binds immediately below the selectivity filter. Pimozide density is shown as a grey surface. Representations are as depicted in Figs. 4*C* and *D*. In *(E)*, the density is semi-transparent. *(F)* Detailed interactions with pimozide. The view is from the side, with one of the two equivalent orientations of pimozide shown. Amino acids that are within van der Waals distance of pimozide are drawn as yellow and tan sticks (corresponding to which subunit they originate from). Dotted lines represent hydrogen bonding distance to the indicated atoms. Helices are depicted by outlined cartoons. K^+^ in S3 and S4 are shown as purple spheres. *(G)* Schematic showing the interactions of pimozide with TWIK-2. Residues contacting pimozide are labeled and depicted as shapes colored blue and green to indicate polar and hydrophobic interactions, respectively. The exposed regions of pimozide (ones that do not have immediate van der Waals contacts) are denoted with grey circles on the chemical structure. This panel was made using Maestro (Schrodinger).

Pimozide makes numerous contacts with the surrounding protein (Figs. 6*F*-*G*). Nitrogen atoms of the piperidine ring and benzimidazole-like moieties are within the hydrogen bonding distance of Thr106 and Thr214, which also coordinate K^+^ at site S4. Numerous van der Waals contacts are made (e.g. with: Val131, Met135, Leu138, Leu246, Met249, Val250, and Leu253). The interactions made with pimozide are more extensive than those for the acyl chains in the apo structure (Figs. 4*E* and 6*F-G*).

The structure suggests that the distribution of ions in the selectivity filter may be altered by the presence of pimozide. In the structure with pimozide, density within the filter is consistent with K^+^ occupancy at sites S3 and S4. Instead of discrete density in S1 and S2, there density located between sites S1 and S2 that could be due to K^+^ (Fig. 6*B*). The polypeptide backbone in the filter and the position of Tyr111 are indistinguishable from the apo structure (Figs. 2*A* and 6*B*), suggesting that the change in density within the filter does not result from protein conformational changes. Rather, the presence of pimozide itself may be responsible for the change. While the apparent change in K^+^ occupancy in the filter is intriguing, caution is warranted in its interpretation because the ∼ 3.2 Å resolution map may not be sufficient to accurately assess ion distribution in the selectivity filter of K^+^ channels (45).

The binding site for pimozide would be accessible from both the intracellular side of the channel (through the pore) and via the fenestrations. Given its highly hydrophobic chemical structure, it seems plausible that pimozide primarily accesses the channel via the hydrophobic core of the lipid membrane through the fenestrations. The positioning of pimozide is close enough to the selectivity filter that it would impede the flow of potassium through the channel (Fig. 6D). We conclude that the mechanism of inhibition is that pimozide blocks the pore.

## Discussion

We report the structure of human TWIK-2. The overall architecture of the channel is similar to that of TWIK-1, but with notable differences. Among these are the positioning of selectivity filter residue Tyr111, additional space between the extracellular cap and the filter, and differences in the positioning of acyl chains within the central cavity.

TWIK-2 is the only K2P channel known to exhibit inactivation (13). ‘C-type’ inactivation in voltage-dependent K^+^ (Kv) channels is thought to involve both protein structural changes within the selectivity filter and reduced ion occupancy (50, 51). In TWIK-2, inactivation depends on the extracellular K^+^ concentration in which low (∼ 5 mM) concentrations of K^+^ stimulate inactivation and high (∼150 mM) concentrations of K^+^ reduce it (13). From comparison with the structures of other potassium channels, the protein backbone of the selectivity filter in the structure of TWIK-2 adopts what appears to be a conductive conformation. Accordingly, in the structure, which was determined using 150 mM K^+^, density is present at all four K^+^ sites within the filter. Inactivation in TWIK-2 is not complete – for instance, even with low (5 mM) external K^+^, currents fall to roughly half of their initial value (13) – therefore, the filter may not experience the marked protein backbone conformational changes observed for certain C-type inactivating K^+^ channels. It is possible that the structure represents a lower conductance state that corresponds to incomplete inactivation in TWIK-2. We speculate that Tyr111 may have a role in the inactivation of TWIK-2 currents. The ‘up’ conformation of Tyr111 observed in the structure is accompanied by a small SF1-P1 pocket behind the upper part of the filter that is normally filled by amino acids in this position in other K^+^ channels. If Tyr111 were to adopt a ‘down’ conformation and fill this pocket, this might rigidify the upper portion of the filter, rendering it more conductive.

Among the K2P channels, TWIK-2 has one of the lowest conductances, which is estimated at 5 pS (13). The structure suggests possible explanations for low conductance. The SF1-P1 pocket and positioning of Tyr111 may make the selectivity filter less conductive than other K^+^ channels. Additionally, the constriction created by Met135 and Val250 within the central cavity could limit ion flow. Accordingly, the narrowest constriction in the central cavity of TWIK-1 is formed by amino acids at corresponding positions (Leu146 and Leu261) and has been proposed to form a ‘hydrophobic cuff’ that limits the rate of ion flow through that channel (52).

Functional data presented here indicates that pimozide inhibits TWIK-2. The structure with pimozide demonstrates that this small molecule binds in the central cavity below the selectivity filter, partially occupies the upper fenestrations in the transmembrane domain, and occludes ion permeation. The binding site overlaps with that observed for acyl chains in the apo structure and is connected to the hydrophobic core of the membrane by the fenestrations. This positioning suggests that pimozide and other such hydrophobic molecules, could access the binding site by first partitioning to the lipid membrane and then diffusing laterally through the fenestrations. The membrane access route suggests that good water solubility may not be a necessary criterion for development of TWIK-2 inhibitors. The walls of the central cavity in which pimozide binds are lined by amino acids that vary among K^+^ channels, suggesting that high-specificity inhibitors could be developed.

The central cavity is a hot spot for the binding of inhibitors of K^+^ channels. Although it doesn’t inhibit TWIK-2 (13), tetraethylammonium (TEA) was one of the first inhibitors shown to occupy this general location (below the selectivity filter) in other K^+^ channels (53, 54). The central cavity has subsequently been shown to be the binding location for other K^+^ channel inhibitors (17, 55). On the basis of mutagenesis and molecular dynamics simulations, NBPA, which inhibits TWIK-2, has been precited to bind in the central cavity at the bottom of the selectivity filter in roughly the location observed for pimozide (41). A recently determined structure of the hERG K^+^ channel in complex with pimozide shows that it binds in an analogous location (56). TWIK-2 has been shown to be activated by polyunsaturated fatty acids (49, 57); the structure suggests that such moieties could bind in the fenestrations adjacent to the central cavity. In addition to the central cavity, K2P channels have other hot spots for pharmacological control (e.g. within the extracellular cap region, in the P1-M4 interface, and within the transmembrane domain) (17). The structure of TWIK-2 suggests that the SF1-P1 pocket behind the filter could be another site for allosteric modulators of channel function. The structural studies presented here may be useful for the design and optimization of TWIK-2 inhibitors with potential utility for controlling the NLRP3 inflammasome.

## Materials and Methods

### Protein expression and purification

cDNA encoding full-length human TWIK-2 (UniProt: Q9Y257) was synthesized (IDT Inc.) and ligated into a mammalian cell expression vector (58) to encode a protein containing a C-terminal GFP tag, which could be removed using PreScission protease. The expression plasmid was transfected into Expi293F cells (Invitrogen, #A14527) for transient expression. Briefly, 1 mg plasmid and 3 mg PEI25k (Polysciences, Inc.) were mixed in 100 ml OptiMEM media (Invitrogen), incubated at room temperature for 20 min, and the mixture was added into 1 L of Expi293 cells (∼3.0 ×10^6^ cells/mL) in Expi293 expression media (Invitrogen). After incubation at 37° C for 16 hours, 10 mM sodium butyrate (Sigma-Aldrich) was added, and the cells were cultured at 30° C for another 48 hours before harvesting.

For protein purification, the cell pellet from 1 L of culture was resuspended in 50 mL lysis buffer [40 mM HEPES pH 7.5, 200 mM KCl, 0.15 mg/mL DNase I (Sigma-Aldrich), 1.5μg/mL leupeptin (Sigma-Aldrich), 1.5 μg/mL pepstatin A (Sigma-Aldrich), 1 mM AEBSF (Gold Biotechnology), 1 mM benzamidine (Sigma-Aldrich), 1 mM PMSF (Acros Organics) and 1:500 dilution of aprotinin (Sigma-Aldrich)], then solubilized by adding 1% (w/v) lauryl maltose neopentyl glycol (LMNG, Anatrace), 0.1% cholesteryl hemisuccinate Tris salt (CHS, Anatrace), and stirred at 4 °C for 1 hr. Solubilized proteins were separated from the insoluble fraction by centrifugation at 60,000 x g at 4° C for 1 hr and the supernatant was filtered through a 0.22-μm polystyrene membrane (Millipore). The TWIK-2-GFP fusion protein was purified by affinity chromatography using a nanobody against GFP (59). The nanobody was coupled to CNBr-activated Sepharose Fast Flow resin (GE healthcare) according to manufacturer’s protocol. 2 mL resin was incubated with the sample with agitation at 4 °C for 1 hr. The beads were washed with 100 mL buffer A (20 mM HEPES pH 7.5, 150 mM KCl, 0.1% LMNG, 0.01% CHS), and purified TWIK-2 protein was eluted by adding PreScission protease (∼0.1 mg; incubating overnight at 4 °C). The sample was further purified by size-exclusion chromatography (SEC, using a Superose 6 increase, 10/300 GL column, GE Healthcare), in SEC buffer (20 mM HEPES, pH 7.5, 150 mM KCl) containing 0.0025% LMNG, 0.00025% CHS, 0.000825% glyco-diosgenin (GDN, Anatrace). The peak fractions were pooled and concentrated to 6 mg/ml using a 100 kDa concentrator and immediately used to prepare cryo-EM grids.

For structure determination of TWIK-2 with 1-[1-[4,4-Bis(4-fluorophenyl)butyl]-4-piperidinyl]-1,3-dihydro-2H-benzimidazol-2-one (pimozide, Tocris), the protein sample was purified in SEC buffer (20 mM HEPES, pH 7.5, 150 mM KCl) containing 0.01% LMNG, 0.001% CHS, 0.0033% glyco-diosgenin (GDN, Anatrace). The peak fractions were pooled and concentrated to 300 uL (to a concentration of ∼ 0.69 mg/mL) using a 100 kDa concentrator. At this point, 1 uL of 150 mM freshly prepared pimozide stock (in DMSO) was added, and the sample was further concentrated to 32 uL and immediately used to prepare cryo-EM grids.

### EM sample preparation and data acquisition

3.5-4 μL of purified samples (TWIK-2 or TWIK-2 with pimozide) were applied to glow-discharged (10s) Quantifoil R 1.2/1.3 grids (Au 400; Electron Microscopy Sciences) and plunge-frozen in liquid nitrogen-cooled liquid ethane, using a Vitrobot Mark IV (FEI), operated at 4 °C, with a blotting time of 3-6 s, using a blot force of ‘0’, and 90% humidity. For the structure of TWIK-2 without pimozide, micrographs were collected using a Titan Krios microscope (Thermo) operated at 300 kV using a Falcon 4i detector (MSKCC) with the Selectris imaging filter (10 eV slit width) in a super-resolution mode. For TWIK-2 with pimozide, micrographs were collected using a Titan Krios microscope (Thermo) operated at 300 kV and equipped with a Falcon 4i detector (NYSBC) with the Selectris imaging filter (20 eV slit width).

### Cryo-EM structure determination

Cryo-EM datasets are summarized in Tables S1; and workflows are summarized in Figs. S2 and S8. For TWIK-2 without pimozide, image processing was performed in RELION 3.1 (60) and cryoSPARC v.4.5.0 (61). Movie stacks were gain-corrected, twofold binned, motion-corrected, and dose-weighted in Relion. Contrast transfer function (CTF) estimates were performed using Patch CTF estimation in cryoSPARC, and micrographs with CTF fit resolution better than 4 Å were selected for further processing. Particles were automatically picked using Template Picker in cryoSPARC. 22,020,573 particles were extracted from 17,758 micrographs in cryoSPARC with a bin factor of 2 and a box size of 384. The particles were subjected to one round of ab initio reconstruction and several rounds of heterogeneous refinements with C1 symmetry. The best class (representing 1,132,393 particles) was selected and extracted without binning (box size of 384). Additional rounds of heterogeneous refinements in cryoSPARC were performed. The best class (representing 866,937 particles) was selected and subjected to two rounds of particle polishing in Relion. The final particles were subjected to 3D classification without image alignment in Relion. The two best classes were subjected to an additional round of ab initio reconstruction. The best class (representing 131,053 particles) gave the final map at 2.85 Å resolution after non-uniform refinement and local refinement with C2 symmetry. Although the final particle set represents a fraction of the initial data, there were no indications throughout the data processing that different conformations of the channel were present in the dataset. The final particle set was selected because it yielded highest resolution structure.

For the structure with pimozide, image processing was performed in RELION 3.1 and cryoSPARC v.3 and v.4. Gain-corrected movies were motion-corrected and dose-weighted in Relion. Contrast transfer function estimates (CTF) were performed with CTFFIND4.1 in Relion (62). Micrographs with CTF fit better than 4 Å were selected for subsequent image processing. Reference-free particle picking was done using the Laplcian-of-Gaussian tool in Relion. 6,263,529 particles were extracted from 12,981 micrographs with a box size of 384 pixels and imported into cryoSPARC. The particles were subjected to one round of ab initio reconstruction and several rounds of heterogeneous refinements (with C1 symmetry). The best class (representing 719,550 particles) was selected and subjected to an additional round of ab initio reconstruction and heterogeneous refinement. From this, 641,558 particles were selected and subjected to four rounds of particle polishing in Relion, which yielded a reconstruction of 3.71 Å after non-uniform refinement with C2 symmetry. The particles were then subjected to 3D classification without image alignment in Relion, followed by non-uniform refinement and local refinement with C2 symmetry, to yield the final map at 3.17 Å resolution. Attempts to computationally separate a discrete conformation of pimozide (e.g. using focused classification with C1 symmetry) were not successful, which is understandable given the location of pimozide on the symmetry axis.

Atomic models were built de novo, refined in real space using COOT (63), and further refined in real-space using PHENIX (64). Structural Figures were prepared with PyMOL (pymol.org) (65), ChimeraX (66), CAVER (67), and HOLE (68).

### Liposome reconstitution and flux assay measurements

Prior to reconstitution, a mixture of lipids was prepared by combining chloroform solutions containing 30 mg of 1-palmitoyl-2-oleoyl-sn-glycero-3-phosphoethanolamine (POPE, Avanti), 10 mg of 1-palmitoyl-2-oleoyl-sn-glycero-3-phosphoglycerol (POPG, Avanti) in a glass tube; evaporating the chloroform using argon; dissolving the lipids in pentane; drying the lipids under argon; and placing the lipids under vacuum overnight while protecting them from light.

Purified water was added to the dried lipids to rehydrate the lipids to a final lipid concentration of 25 mg/ml. The lipids were incubated with end-over-end mixing for 1 hour, and the mixture was sonicated in a water bath for a total of 5 min in 1-min on/off intervals. The lipids were subsequently destabilized by adding detergent (C12E10) and buffer and allowed to rotate at room temperature for 1 hour with end over end mixing. The final solution contained 10 mM HEPES-NMDG pH 7.6, 150 mM KCl, 1% polyoxyethylene(10)dodecyl ether • 3,6,9,12,15,18,24,27,30-decaoxadotetracontan-1-ol (C12E10, Anatrace), and 20 mg/ml of the lipid mixture.

For reconstitution into liposomes, freshly prepared protein in SEC buffer (20 mM HEPES-NMDG, pH 7.6, 150 mM KCl, 0.001% LMNG, 0.0001% CHS, and 0.00033% GDN) was concentrated to ∼1 mg/ml using a 50-kDa MWCO concentrator (Amicon Ultra-2) and mixed 1:1 with the C12E10-solubilized lipids to yield the desired protein-to-lipid ratio (1:100, for example, 30 μg of protein would be mixed with 3 mg of PE:PG). After mixing end-over-end for one hour at 4°C, detergent was removed by dialysis (50,000 Da molecular weight cutoff) at 4°C against a total of 10 liters of buffer (20 mM HEPES-NMDG pH 7.0, 150 mM KCl) with daily buffer exchanges over the course of 5 days. On days 2 through 5, the daily buffer exchange included activated Bio-Beads SM2 resin (Bio-Rad) in the buffer at a ratio of ∼ 1 g of beads (wet weight) per 2L of buffer. Empty liposomes (without protein) were prepared in parallel in the same manner using purification buffer instead of purified protein. Following dialysis, the liposomes were sonicated for approximately 20 s in a water bath, divided into 50 uL aliquots, and flash-frozen in liquid nitrogen for storage at −80°C.

The flux assay was based on previously published methods (22, 69). Vesicles were thawed in a 37°C water bath, sonicated (for ∼20 s, in 10-s intervals), and incubated at room temperature for 2 hours. The flux assay buffer consisted of 4 mM HEPES-NMDG (pH 7.0), bovine serum albumin (0.5 mg/ml), 2 μM 9-amino-6-chloro-2-methoxyacridine [ACMA; Sigma-Aldrich, from a 2 mM stock solution in dimethyl sulfoxide (DMSO)], and 125 mM NaCl. For inhibitor studies (with pimozide, ML365, or BaCl_2_), the compounds were included in the flux assay buffer at the indicated concentrations. Data were collected on a SpectraMax M5 fluorometer (Molecular Devices) using the SoftMax Pro 7 software package. Fluorescence intensity measurements were collected every 30 s with excitation and emission wavelengths of 410 and 490 nm, respectively.

The course of the experiment was as follows: The fluorescence of the flux assay buffer (1 ml) was recorded for 120 s, liposomes were added by 500-fold dilution, and 1 μM of the proton ionophore carbonyl cyanide m-chlorophenyl hydrazone (CCCP; Sigma-Aldrich; from a 1 mM stock solution in DMSO) was added at the 300-s time point. The sample was gently mixed with a pipette after each addition. To confirm that the vesicles were intact and to establish as minimum fluorescence baseline, 2 nM valinomycin (Sigma-Aldrich; from a 2 μM stock solution in DMSO) was added after the 1260-s time point (valinomycin allows selective K^+^ efflux from the vesicles, creating a negative charge within them that is used to drive the uptake of H^+^).

## Author Contributions

N.K.K. and C.W. performed cryo-EM and related studies. B.D.D. performed functional experiments. S.B.L. directed the research. All authors contributed to data analysis and the preparation of the manuscript.

## Competing Interests Statement

The authors declare that they have no competing interests. This work was performed before B.D.D. became an employee of Genentech Inc.

## Acknowledgements

We thank members of the S.B.L. laboratory for helpful discussions and M. J. de la Cruz for assistance with cryo-EM at MSKCC. This work was supported by NIH grant R35GM131921 (to S.B.L.), NIH core facilities grant to MSKCC (P30CA008748), and NIGMS F31 fellowship 1F31GM143882 (to B.D.D.). Some of this work was performed at the National Center for CryoEM Access and Training (NCCAT) and the Simons Electron Microscopy Center located at the New York Structural Biology Center, supported by the NIH Common Fund Transformative High Resolution Cryo-Electron Microscopy program (U24 GM129539, and NIGMS R24 GM154192) and by grants from the Simons Foundation (SF349247) and NY State Assembly.

## Data and materials availability

Atomic coordinates and maps of human TWIK-2 and human TWIK-2 in complex with pimozide have been deposited in the PDB (ID: 9MEK and 9MEL, respectively) and EMDB (EMD-48216 and EMD-48217, respectively). All data needed to evaluate the conclusions in the paper are present in the paper and/or the Supplementary Materials.

## Supporting Information

**Fig. S1.**
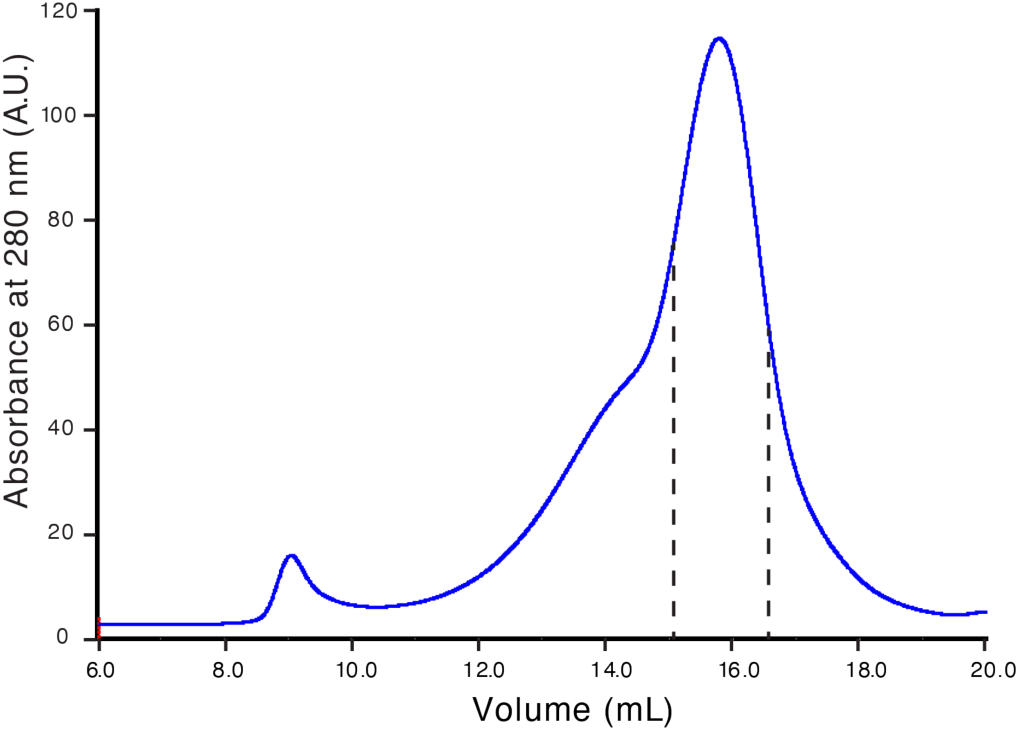
Size-exclusion chromatography trace of purified TWIK-2. The region used to prepare cryo-EM grids is indicated by dashed lines.

**Fig. S2.**
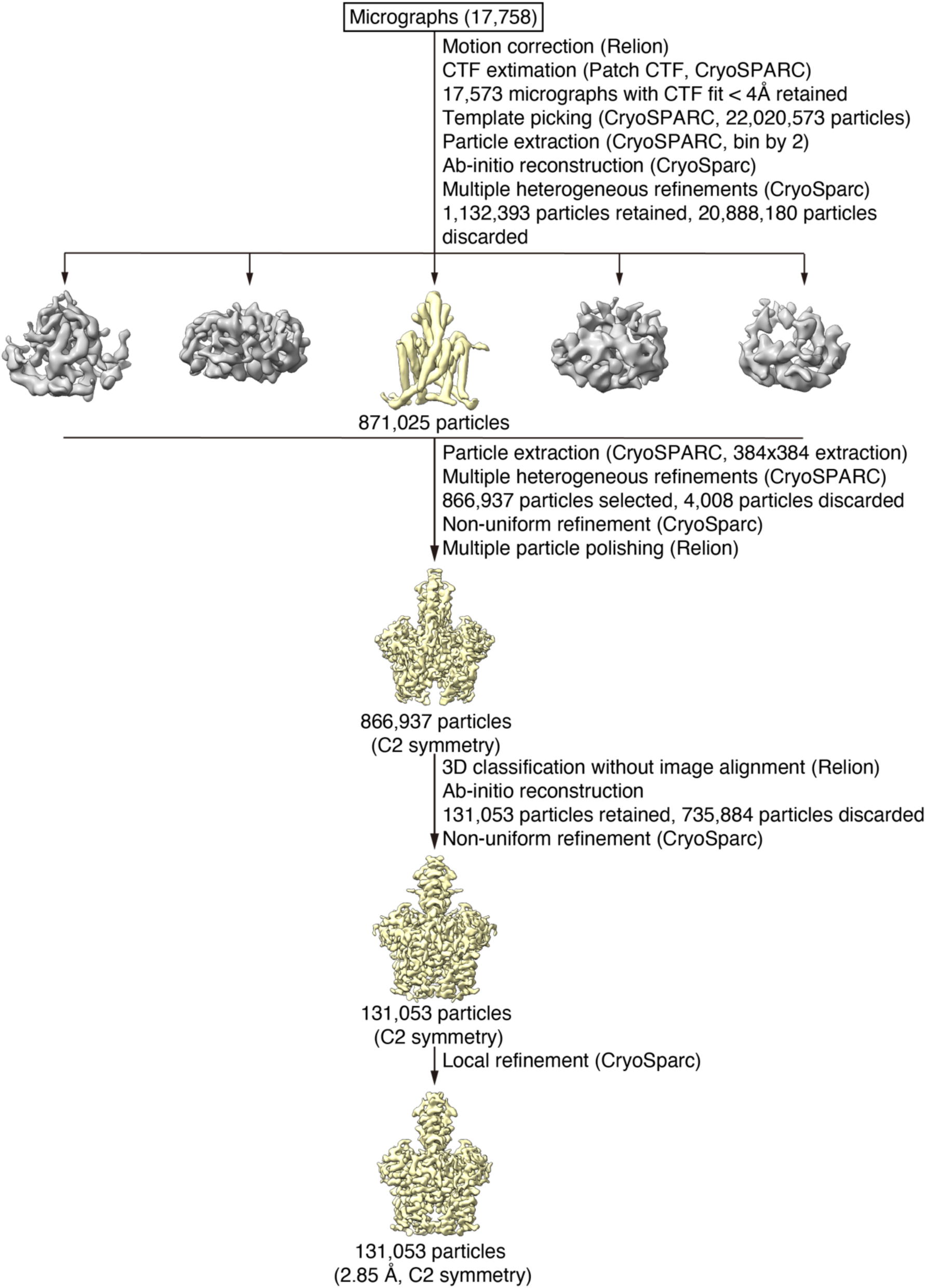
The cryo-EM data processing workflow for TWIK-2 (without pimozide). Details are described in Materials and Methods.

**Fig. S3.**
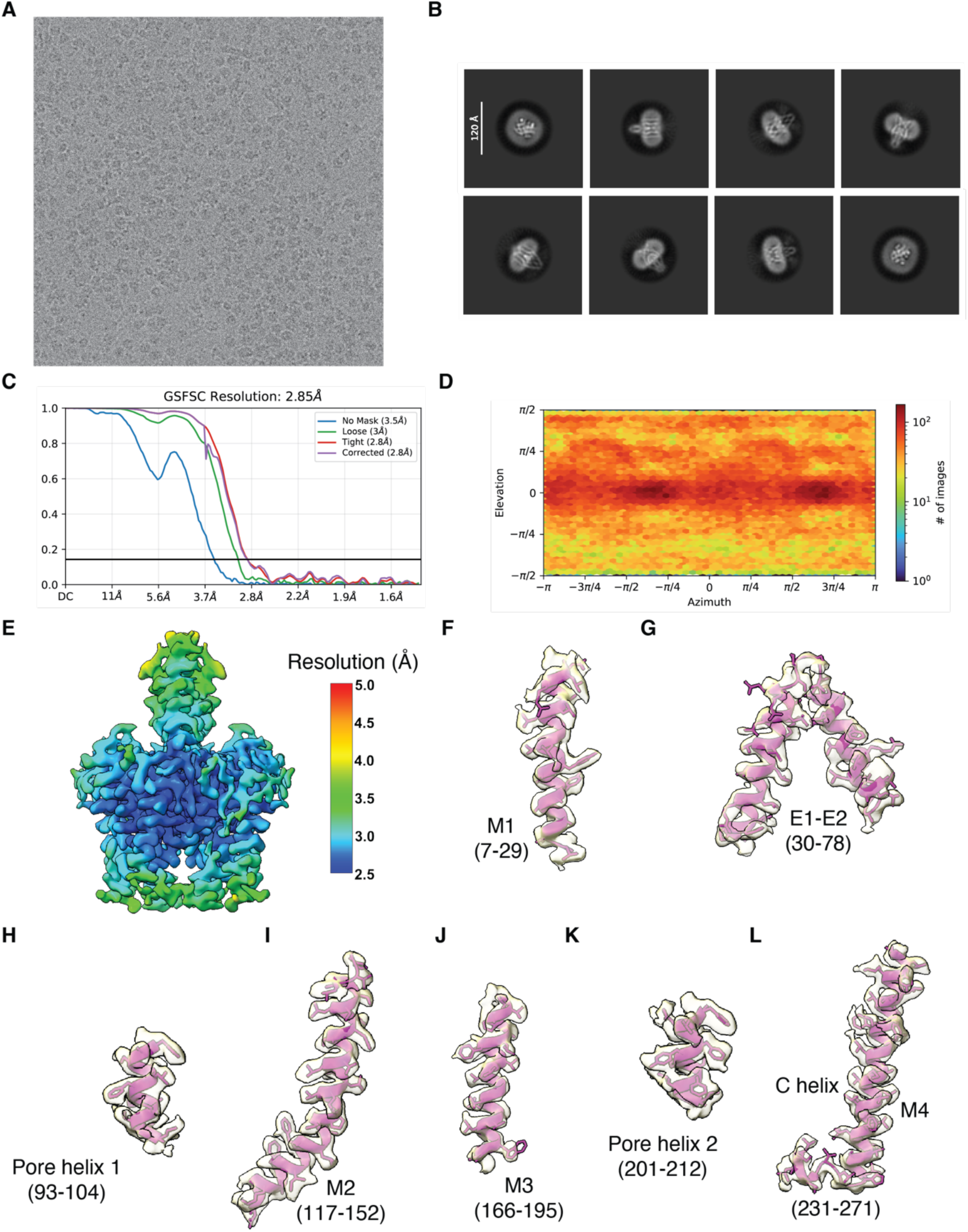
Micrograph, 2D class averages, and analysis of the cryo-EM map of TWIK-2. *(A)* Representative raw image. *(B)* Representative 2D class averages. *(C)* Half-map FSC curves. *(D)* Angular distribution of particles used in the final reconstruction. *(E)* Estimations of local resolution. *(F-L)* Densities (semi-transparent surface rendering) for indicated regions are shown in the context of the atomic model (cartoons and sticks). Details are described in Materials and Methods.

**Fig. S4:**
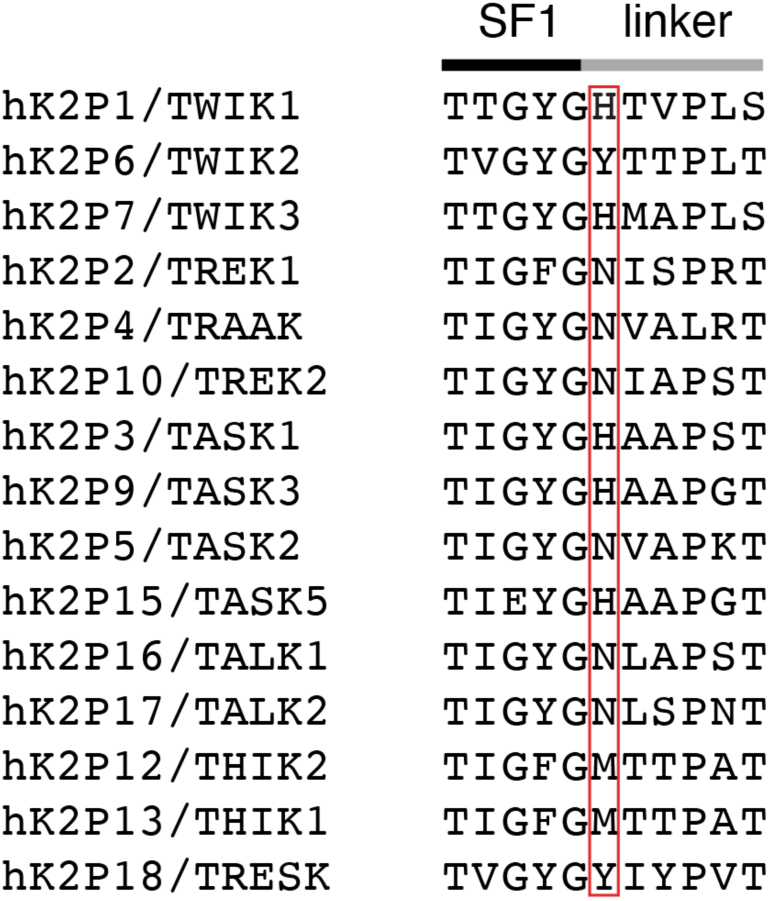
Sequence alignment of the SF1 and SF1-M2 linker region among all fifteen human K2P channels. The boxed region corresponds to Tyr111 of TWIK-2.

**Fig. S5.**
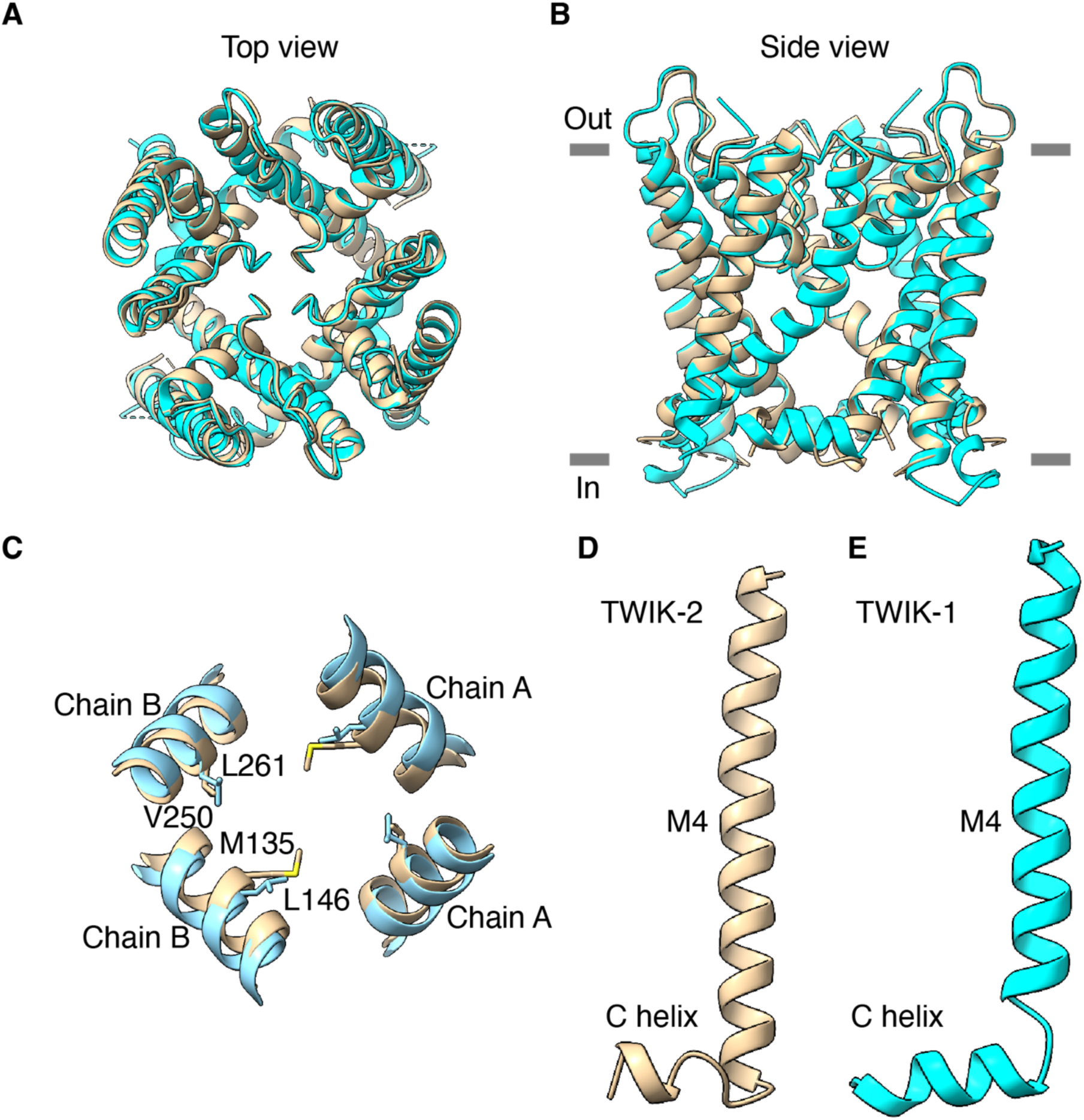
Comparison of the transmembrane regions of TWIK-1 and TWIK-2. *(A-B)* Overall views, showing cartoon representations of TWIK-1 (cyan, PDB ID 3UKM) and TWIK-2 (tan). *(C)* The hydrophobic cuff in the central cavity, showing Met135 and Val250 from TWIK-2 and corresponding residues (Leu146 and Leu261) of TWIK-1. *(D-E)* The M4 and C helices of TWIK-2 and TWIK-1.

**Fig. S6.**
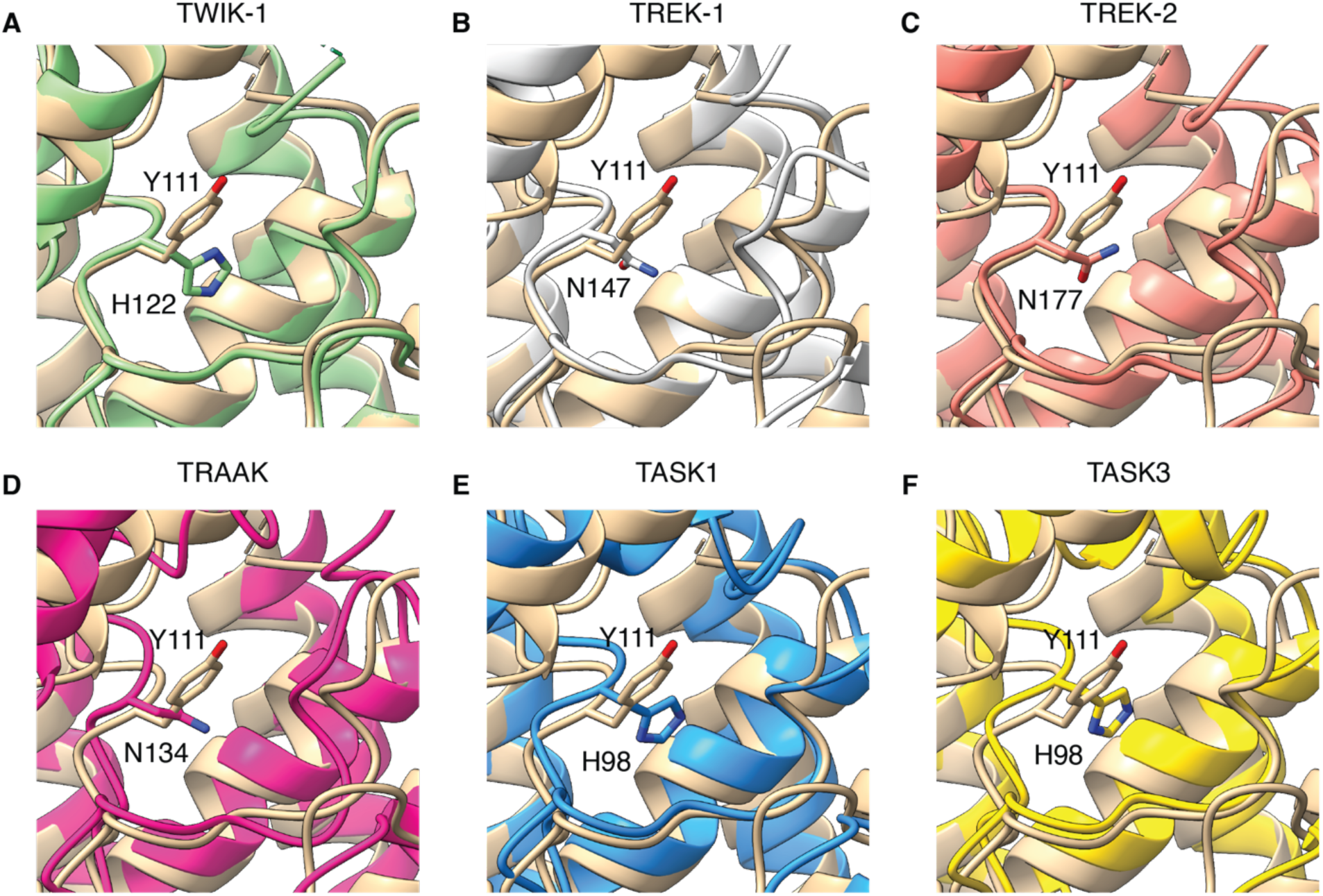
Positioning of Tyr111 in TWIK-2 compared with corresponding residues of other K2P channels. The view is analogous to Fig. 2F. TWIK-2 is shown in tan in each panel and compared with: *(A)* His122 of TWIK-1 (green, 3UKM), *(B)* Asn147 of TREK1 (grey, 8DE8), *(C)* Asn177 of TREK2 (orange, 4BW5), *(D)* Asn134 of TRAAK (violet, 3UM7), *(E)* His98 of TASK1 (blue, 6RV4), and *(F)* His98 of TASK3 (yellow, 8K1J).

**Fig. S7.**
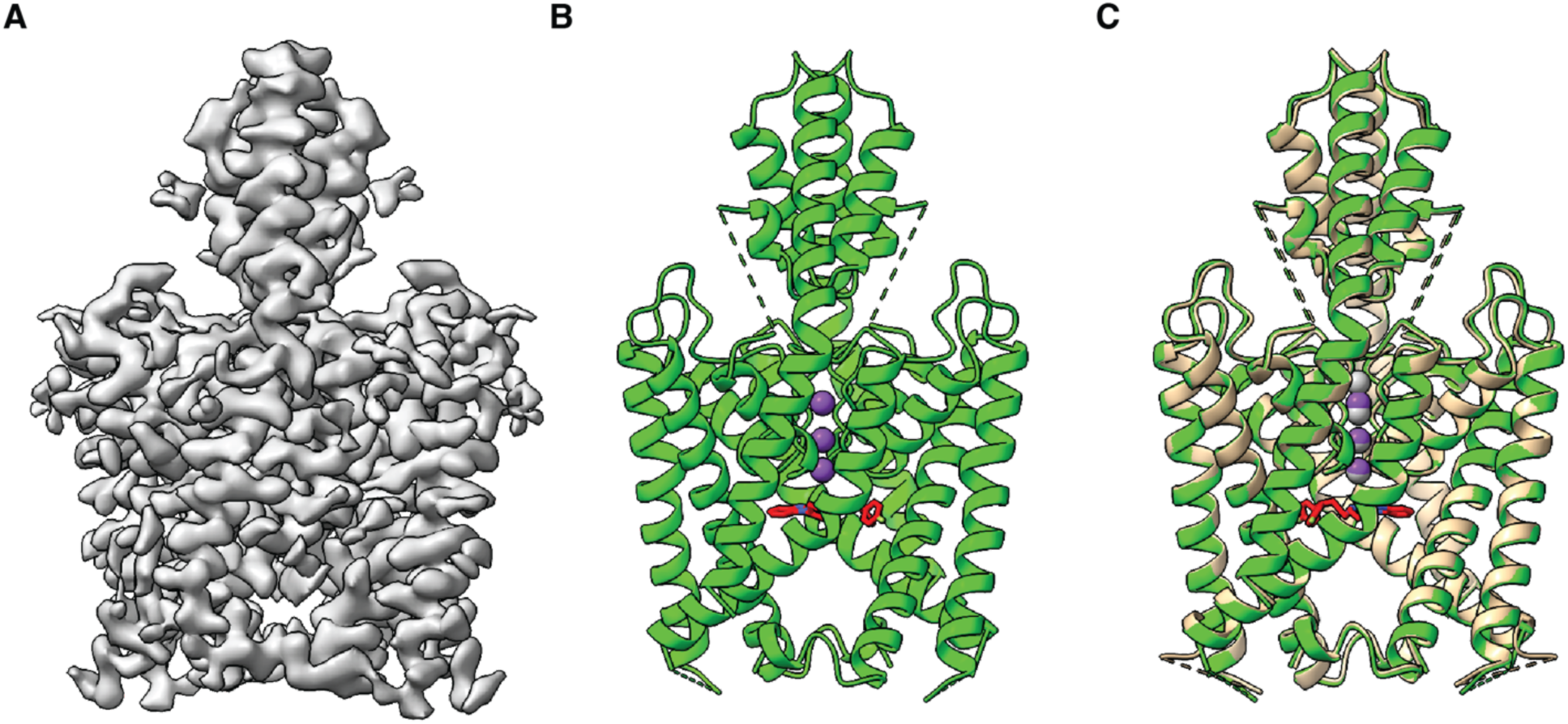
Overall structure of TWIK-2 in complex with pimozide. *(A)* Cryo-EM map of the complex. *(B)* Atomic model, shown in cartoon representation with K^+^ as purple spheres and pimozide as magenta sticks. Dashed lines indicate disordered loop regions. *(C)* Superposition of the overall structures without (tan) and with pimozide (green). Potassium ions in the structure without pimozide are depicted as grey spheres.

**Fig. S8.**
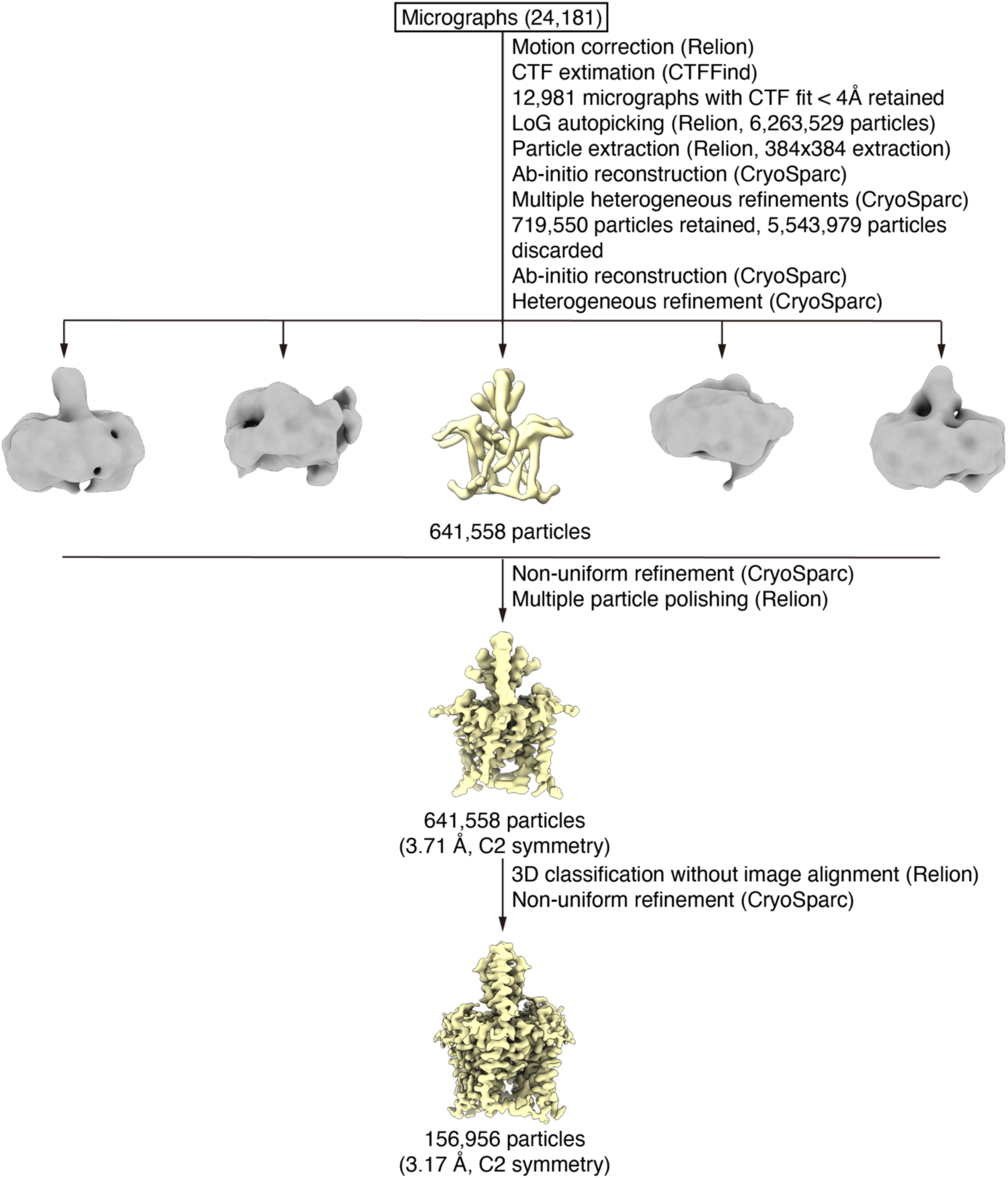
The cryo-EM data processing workflow for the complex of TWIK-2 with pimozide. Details are described in Materials and Methods.

**Fig. S9.**
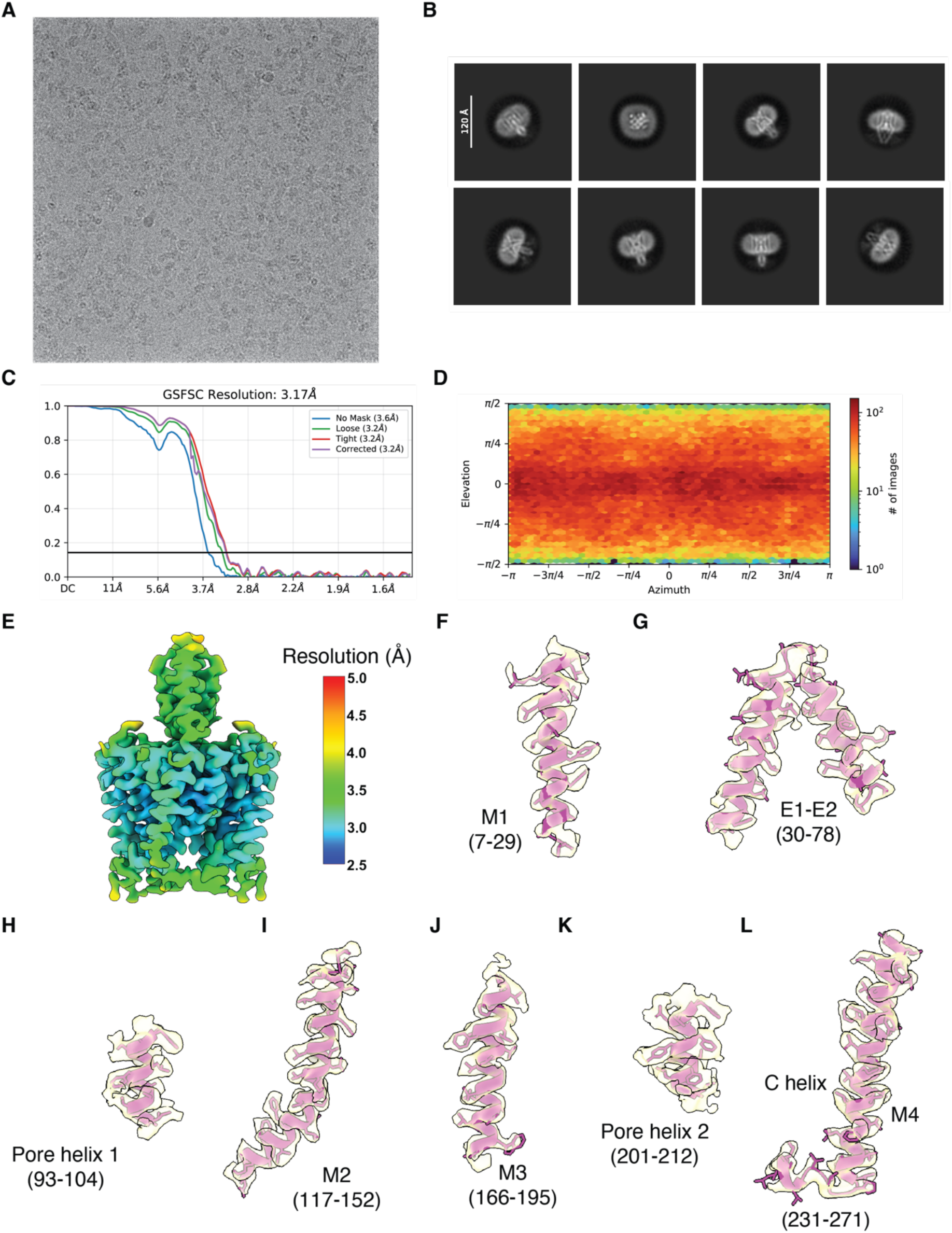
Micrograph, 2D class averages, and analysis of the cryo-EM map of TWIK-2 with pimozide. *(A)* Representative raw image. *(B)* Representative 2D class averages. *(C)* Half-map FSC curves. *(D)* Angular distribution of particles used in the final reconstruction. *(E)* Estimations of local resolution. *(F-L)* Densities (semi-transparent surface rendering) for indicated regions are shown in the context of the atomic model (cartoons and sticks). Details are described in Materials and Methods.

**Fig. S10.**
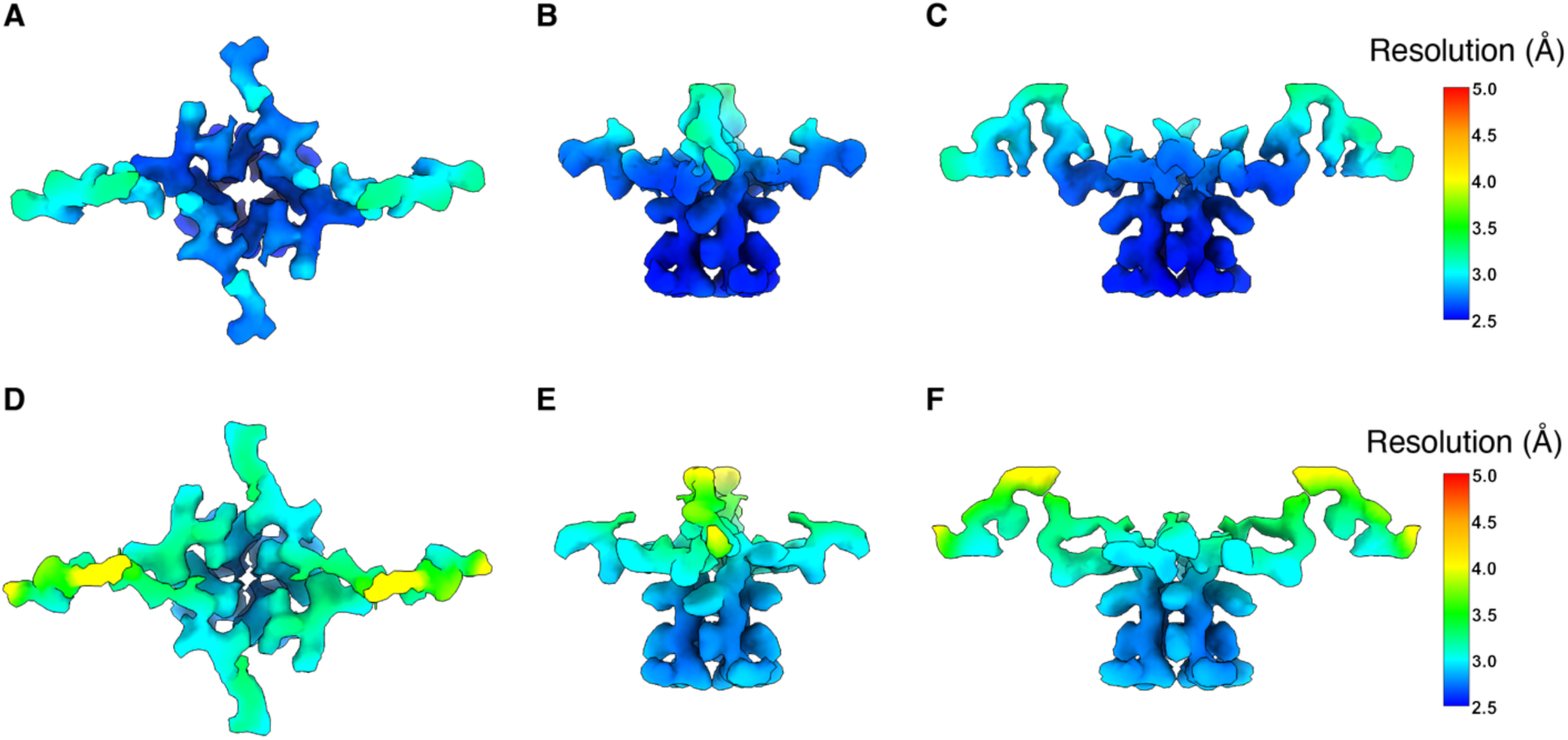
Local resolution estimations of selectivity filter and linker regions for complexes without *(A-C)* and with pimozide *(D-F)*. *(A)* Top view. *(B-C)* orthogonal side views. *(D-F)* Analogous views for the complex with pimozide. Densities are depicted as surfaces, colored according to local resolution, and shown only for the selectivity filter and linker regions.

**Fig. S11.**
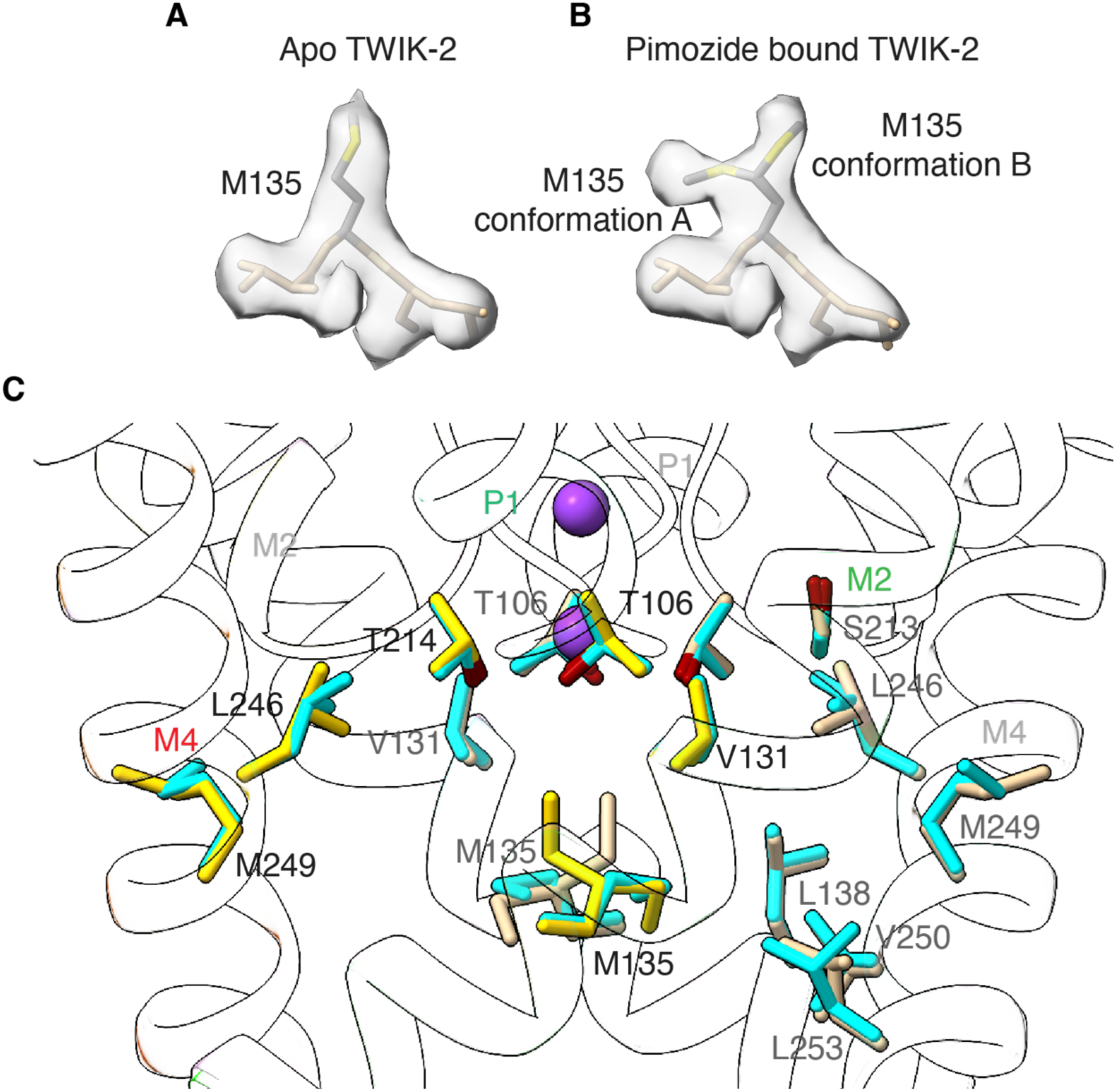
Comparison of amino acid positions with and without pimozide. *(A-B)* Conformation of Met135 without *(A)* and with pimozide *(B)*. Density is shown as a semi-transparent surface for Met135 and flanking residues (sticks). The alternate conformations of Met135 with pimozide are indicated. *(C)* Conformations of amino acids with and without pimozide. The side chains of amino acids that contact pimozide are drawn as sticks. Conformations without pimozide are shown in cyan; those in the structure with pimozide are tan (one subunit) and yellow (the other subunit).

**Table S1.** Data collection, model refinement, and validation statistics.

|  | TWIK2-apo<br>(PDB: 9MEK, EMD-48216) | TWIK2-pimozide<br>(PDB: 9MEL, EMD-48217) |
| --- | --- | --- |
| Data collection and processing |  |  |
| Microscope | FEI Titan Krios (MSKCC) | FEI Titan Krios (NYSBC) |
| Camera | Falcon 4i | Falcon 4i |
| Magnification | 165,000× | 165,000× |
| Voltage (kV) | 300 | 300 |
| Electron exposure (e<sup>-</sup>/Å<sup>2</sup>) | 60.76 | 51.38 |
| Defocus range (μm) | -1.0 ~ -2.0 | -1.0 ~ -2.5 |
| Pixel size (Å) | 0.725 | 0.725 |
| Software | RELION 3.1, cryoSPARC v4.5.0 | RELION 3.1, cryoSPARC v4.5.0 |
| Symmetry imposed | C2 | C2 |
| Initial particle images (no.) | 22,020,573 | 6,263,529 |
| Final particle images (no.) | 131,053 | 156,956 |
| Overall map resolution (Å) | 2.85 | 3.17 |
| FSC threshold 0.143 |  |  |
| Local map resolution range (Å) | 2.5-4.0 | 2.7-4.2 |
| Atomic model refinement |  |  |
| Software | Phenix 1.21 real-space-refine | Phenix 1.21 real-space-refine |
| Initial model (PDB code) | N/A | N/A |
| Model resolution (Å) | 2.8 | 3.1 |
| FSC threshold 0.5 |  |  |
| Map sharpening B factor (Å<sup>2</sup>) | - 89.3 | -124.2 |
| Model composition |  |  |
| Non-hydrogen atoms | 3,594 | 3,589 |
| Protein residues | 478 | 476 |
| Ligands | 4 K<sup>+</sup>, 2 acyl | 2 K<sup>+</sup>, 1 pimozide |
| Water | 0 | 0 |
| Average B factors (Å<sup>2</sup>) |  |  |
| Protein | 64.48 | 80.08 |
| Ligands | 39.81 | 44.37 |
| R.m.s. deviations from ideality | 0.004 | 0.004 |
| Bond lengths (Å) | 0.910 | 0.668 |
| Bond angles (°) |  |  |
| Validation |  |  |
| MolProbity score | 1.58 | 1.69 |
| Clashscore | 6.85 | 4.72 |
| Poor rotamers (%) | 0.00 | 0.00 |
| Ramachandran plot |  |  |
| Favored (%) | 97.85 | 96.12 |
| Allowed (%) | 2.15 | 3.88 |
| Disallowed (%) | 0.00 | 0.00 |

## Notes

### Competing Interest Statement

The authors have declared no competing interest.

